# Evaluation of Trypanosoma brucei Phosphofructokinase Allosteric Inhibition: An *In-Silico* Study

**DOI:** 10.64898/2026.06.16.732740

**Authors:** Gediminas Gumbis, Douglas R. Houston

## Abstract

Human African trypanosomiasis, caused by a protozoan parasite *Trypanosoma brucei*, is a neglected tropical disease for which well-tolerated, conveniently administered, and highly efficacious medicines are still missing. Previously, *T. brucei* Phosphofructokinase was targeted by small-molecule inhibitor development efforts. This approach has shown promise both *in vitro* and *in vivo*. In this study, we have used these wet-lab results, evaluated the compounds already characterised by Molecular Dynamics simulations, found relationships between *in silico* and wet-lab data and used these observations to evaluate compounds that we selected through several different approaches of virtual screens. We observed that inhibitor-ATP interactions are highly predictive of the inhibitory activity. Several compounds selected through virtual screens have outperformed previously characterised compounds.

## INTRODUCTION

Human African trypanosomiasis (HAT), also called ‘sleeping sickness’, is caused by a protozoan parasite *Trypanosoma brucei* [1,2]. *T. brucei* is transmitted by bites of infected tsetse flies (*Glossina*). Two forms of *T. brucei* have been identified as causing two different forms of disease – slow-progressing *T. brucei gambiense,* found in western and central Africa, and more rapidly progressing *T. brucei rhodesiense*, found in eastern and southern Africa [2]. Trypanosomes causing animal trypanosomiasis include *T. brucei, T. evansi, T. equiperdum, T. vivax, and T. congolense* [3]. HAT has two stages – during the first one, the parasites are deposited in the skin, escape into the blood and lymphatics, where they circulate, thus called the haemolymphatic stage. During the second stage, also called meningoencephalitic, there is central nervous system (CNS) involvement [2].

During the haemolymphatic stage, the parasite has evolved to rely on host blood glucose as a predominant source of energy. It was shown that during this stage of its life-cycle, *T. brucei* depends on glycolysis for its ATP production [4]. Previous studies have, therefore, tested the potential of this pathway to be used as a target for drug development efforts. More specifically, inhibition by small-molecule compounds and RNA interference-mediated knockdowns of *T. brucei* Phosphofructokinase (TbPFK) led to the killing of the pathogen [5,6].

Across the tree of life, Phosphofructokinases (PFKs) are distributed into two non-homologous superfamilies in terms of their amino acid sequence similarities: Rossmann fold-containing PFK Superfamily (also referred to as PFKA) and members of the Ribokinase-Like Superfamily of sugar kinases (also referred to as PFKB) [7,8]. The PFKB family also includes other kinases, not only those that phosphorylate fructose 6-phosphate [9]. Independent from this classification, PFKs also differ in terms of phosphate-donor utilised, which can be ATP, ADP or inorganic pyrophosphate (PP_i_) [7]. Many PFKs exhibit two functionally distinct states: functioning, high-affinity R state and inhibited T state, which were first established to exist in *G. stearothermophilus* [10]. These states were also shown to exist in TbPFK structures acquired through X-ray crystallography [11]. Comparison of T and R state structures of TbPFK have identified that the T-to-R transition involves large movements of catalytic Asp229 and Asp231 residues [6].

Regarding the disease vector, *T. brucei gambiense* is mainly transmitted to humans by tsetse flies of the *Glossina palpalis* group, the main reservoir of disease being humans, although pigs and some wild animals could also play a smaller role as disease reservoirs [12]. *T. brucei rhodesiense* infections are transmitted by tsetse flies of the *Glossina morsitans* group, in this case, humans, ungulate wildlife (mainly antelopes) and cattle act as reservoirs [12].

HAT is endemic to sub-Saharan Africa [13]. According to the WHO, the incidence of HAT differs according to the countries and regions considered. According to the available data for 2018-2022:

- 1 country (the DRC) reported 61% of the total cases.
- 8 countries declared 10–100 cases.
- 7 countries reported 1–10 cases.
- 5 countries identified only sporadic cases in the last decade.
- 15 countries have not reported cases for over a decade. Transmission of the disease seems to have stopped in some of these countries, but this is not fully assessed yet [13].

There are six medications used to treat HAT: fexinidazole, pentamidine, nifurtimox (used in nifurtimox-eflornithine combination therapy), eflornithine (can be used as monotherapy as well), suramin, and melarsoprol. All of these medicines have a number of shortcomings, including varying levels of side effects, which can be severe, such as in the case of melarsoprol, which, due to its toxicity, is now reserved for second-stage rhodesiense HAT patients for whom there are no other medication choices available. A need for new HAT medicines which would have higher efficacy, more convenient administration routes and more favourable side effect profiles is often stressed [14,15]

In this study, we have run molecular dynamics (MD) simulations of 11 compounds previously characterised by McNae and colleagues, supplemented this with several thorough *in silico* screening protocols involving docking, molecular clustering, pharmacophore modelling, and semi-empirical quantum mechanics simulations. In total, we ran up to 500 ns of MD simulations for 35 small molecules together with TbPFK. Our aim was to utilise *in silico* approaches to expand the capabilities of wet-lab approaches used before: 1) to try and find additional compounds with similar or better performance, 2) to add to the understanding of the mechanism of the inhibition of TbPFK.

## MATERIALS AND METHODS

### Preparation of structures

Two protein systems were prepared: one with ATP and ligand, and apoprotein. For this, we have used SWISS-MODEL [16], PEP-FOLD3 [17], and PyMOL. The one with ATP and ligand was built accordingly: residues 1-9 are modelled using PEP-FOLD3; 10-18 are from a homology model made with 6sy7 as a template; 19-330 derive from crystal structure (6QU3); 331-340 are from a homology model made with 6QU3 as a template; 341-486 derive from crystal structure (6QU3), and 487 was added using PyMOL.

The apoprotein system was built accordingly: 1-8 were modelled using PEP-FOLD3; 9-66 derive from crystal structure (2HIG); 67-74 were taken from homology model made with 6QU3; 75-332 derive from crystal structure (2HIG); 333-336 were taken from homology model with 6QU3; 337-454 were taken from a crystal structure (2HIG); 455-486 were derived from homology model with 6QU3; and 487 was added with PyMOL.

Subsequently, Hydrogen atoms were added using PyMOL to both of these systems.

In terms of preparation of CTCB ligand structures, CTCB-360 and CTCB-405 were taken from crystal structures, while others were built in PyMOL using these structures as a base.

### Life Chemicals kinase inhibitor library docking

Three initial receptor structures were used for docking: 1) 6QU3 crystal structure, 2) structure derived from MOPAC (GNORM=5) run of 6QU3 apoprotein, 3) structure derived from MOPAC (GNORM=5) run of 6QU3 apoprotein+CTCB360+ATP+Mg^2+^.

During protein preparation, all water molecules and non-standard heteroatoms were removed to prevent steric clashes and docking artifacts. Missing atoms were reinserted into optimized positions using PDB2PQR [18,19]. Protonation states were predicted at pH=7.4 using the PropKa 3 algorithm [20,21]. Atomic charges and radii were assigned based on the AMBER f99 force field. Final structures were converted to PDBQT format using the prepare_receptor4.py script from AutoDockTools 1.5.7 [22].

For small molecule ligand preparation, hydrogen atoms were assigned according to the predicted protonation state of functional groups at pH=7.4 using OpenBabel 3.1.1 [23]. To enable conformational flexibility during docking, rotatable bonds were defined and Gasteiger charges were assigned using the prepare_ligand4.py utility.

Docking was conducted using AutoDock Vina (will be henceforth referred to as Vina) and AutoDock-GPU (will be henceforth referred to as AutoDock). The search space for both docking engines was defined as a 8 Å cubic box centered on the center of gravity of the native ligand (CTCB360). To maximize the probability of identifying the global energy minimum, the exhaustiveness parameter for Vina was increased to 100. Grid maps for AutoDock were pre-calculated using AutoGrid with a spacing of 0.2 Å. This refined grid resolution was chosen to improve docking accuracy over the standard 0.375 Å spacing, as recommended by Houston and Walkinshaw (2013).

For the docking simulations, we utilized AutoDock to leverage parallel processing. The following search parameters were implemented: The population size was set to 300, with the number of GA iterations increased to 30,000 per docking run, a maximum of 10,000,000 evaluations was permitted per ligand to ensure convergence, and to avoid starting-point bias, initial translation, rotation, and torsion angles were randomized for each docking attempt.

### Enamine library docking

Docking of the enamine library followed the same protocol as that outlined for Life Chemicals kinase inhibitor library. However, this library was only docked against 6QU3 crystal structure. This choice was made because during previous docking of Life Chemicals kinase inhibitor library only the compounds docked against the 6QU3 crystal structure were found to have negative binding energies to the protein and this interaction was hypothesised to be crucial for the inhibitory effect of ligand.

### SCORCH analysis and Hit selection

Following initial docking, the structural poses generated via Vina and AutoDock were filtered using the SCORCH pipeline to isolate high-confidence hits and minimize false positives [24].

Afterwards, the compounds were subjected to a multi-tiered filtering cascade: individual compounds were only retained if they achieved a Vina SCORCH score greater than 0.3, a Vina SCORCH certainty greater than 0.5, and an AutoDock SCORCH score greater than 0.1. For the subset of molecules successfully satisfying all three criteria, a consensus binding score was calculated by averaging the best scores acquired by Vina and AutoDock. Compounds were ranked based on this score, and top three compounds from each screen were retained for further analyses.

### Semi-empirical quantum mechanics simulations

Performed using MOPAC in several stages, evaluating progress after each step [25]. PM6-D3H4 and MOZYME were used, EPS was set to 78.4 [26,27]. First, only Hydrogen atoms were optimised, afterwards, GNORM=10 and GNORM=5 were used to optimise the entire protein-ligand complexes.

### Compound parametrisation for molecular dynamics simulations

Initial topology files and force field parameters for all small organic molecules were generated using SwissParam [28]. Force field parametres are exctracted from the Merck Molecular Force Field (MMFF). However, Van der Waals parameters are assigned based on the closest matching atom types within the CHARMM22 force field. Prior to submission to the parameterization server, all molecules were converted to the .mol2 format, verifying the inclusion of explicit 3D spatial coordinates and all necessary hydrogen atoms.

### Molecular dynamics simulations

Using VMD, protein-ligand complexes were made, solvated (13 Å in each direction), and, subsequently, the solvation box was ionised with 0.15 M of Na^+^ and Cl^-^ [29]. Molecular Dynamics (MD) simulations followed; for this, we utilised NAMD 2.14 [30]. CHARMM36m was used for the protein atoms. Minimisation was carried out for 20,000 steps, while a timestep of 2 fs. Following minimisation, the systems were equilibrated for 1 ns, and, finally, longer MD simulations were carried out. During equilibration and production stages, 310 K constant temperature was maintained. Initially, 100 ns of production simulation was run, but for promising compounds, depending on ligand position in the binding pocket and binding free energy estimates, the production runs were continued up to 300 ns or 500 ns. CaFE was utilised to perform molecular mechanics Poisson-Boltzmann surface area (MM/PBSA) calculations which produced binding affinity estimates [31]. During CaFE runs, NAMD was used for the MM component, APBS for PB calculations, and VMD for SA calculations [32]. All CaFE runs used default parameters.

### Pharmacophore modelling

Pharmacophore modelling was conducted using LigandScout [33]. We used compounds from Life Chemicals kinase inhibitor library and the CTCB ligand set that were subset from all analysed compounds to produce the pharmacophore. The software produced ten candidate pharmacophore models, of which, the one with the highest performance was chosen.

### Molecular Clustering

To evaluate the structural diversity of the screening hits and ensure broad scaffold representation, the top 100 performing compounds isolated from the Enamine high-throughput virtual screen were subjected to molecular clustering using ChemBioServer [34]. Distance method selected: Soergel (Tanimoto Coefficient), clustering method: Ward’s linkage, clustering threshold: 1 [35]. We then looked at how the best 10 compounds are distributed amongst the clusters formed.

These 10 best compounds were to be analysed further. Beyond these, we would also take the best compounds from the clusters that would perform similarly to clusters, in which the best 10 compounds were located, in terms of average affinity.

### ADME prediction

We have used SwissADME for the absorption, distribution, metabolism, and excretion (ADME) prediction [36]. A range of predicted variables were extracted. The structural descriptors calculated for each compound included molecular weight, consensus lipophilicity, topological polar surface area, as well as the count of hydrogen bond donors and acceptors. Drug-likeness evaluations were benchmarked against five established rule-based filtering variants: Lipinski (Rule of Five), Ghose, Veber, Egan, and Muegge. Gastrointestinal (GI) absorption and blood-brain barrier (BBB) permeation capabilities were evaluated to anticipate systemic distribution. Finally, structural liabilities were flagged using the Pan-Assay Interference Compounds (PAINS) and Brenk structural alert filters, while overall synthesis feasibility was assessed via the fractional synthetic accessibility (SA) score.

### Isoelectric point and charge prediction

Theoretical isoelectric points (pI) and pH-dependent molecular net charges were calculated using Chemicalize web platform (ChemAxon, https://chemicalize.com/). Calculations were executed during February and March of 2026. Net charges were specifically evaluated at a physiological benchmarking value of pH = 7.4 to guide the assessment of EC_50_ values.

## RESULTS

### Compounds analysed

Here we give IUPAC names generated by Chemicalize, structures, and SMILES codes generated by https://sciencecodons.com/tools/pdb-to-smiles-converter/. The IUPAC names and structures can be found Supplementary Table 1, while SMILES codes are reported in Supplementary Table 2.

### Life Chemicals kinase inhibitor library docking results

Here we report the results of Life Chemicals kinase inhibitor library docking (Table 1).

**Table 1:**
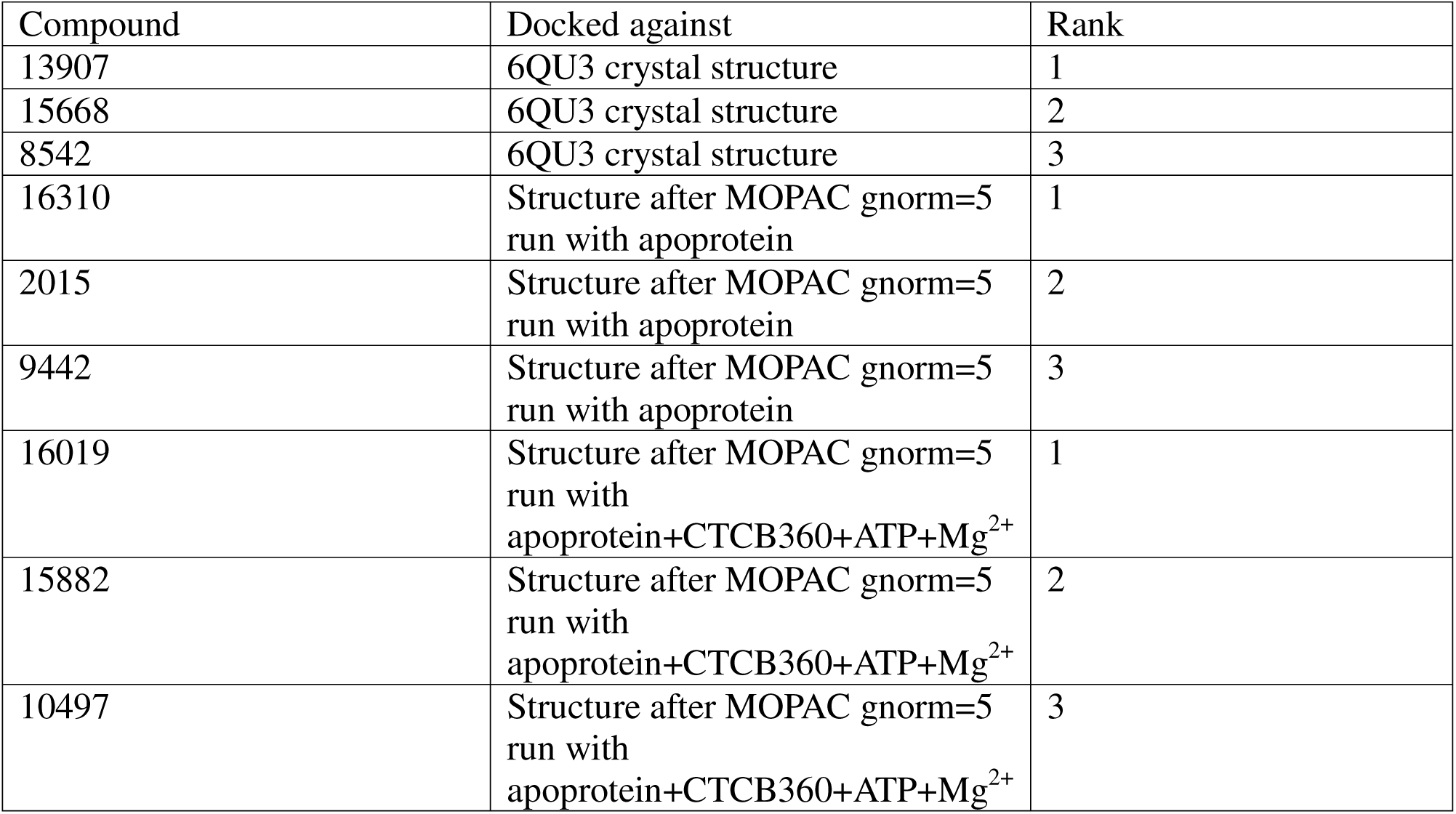
Selected Life Chemicals kinase inhibitor library compounds.

### MM-PBSA results

We have run up to 500 ns of MD simulations for the studied compounds. Binding energies were calculated at two time points: at 100 ns, taking into account time from 0 ns to 100 ns, and at 500 ns, taking into account time from 300 ns to 500 ns. For each compound and time point, at which estimates were made, binding energies of ligand-apoprotein, ligand-ATP, and ligand-apoprotein-ATP complex were made. At 300 ns, binding energy estimates were not made, but rather, it was visually evaluated whether the ligand stayed in the binding pocket or escaped it. Results are reported in Supplementary Tables Supplementary Table 3, Supplementary Table 4, Supplementary Table 5, and Supplementary Table 6 for CTCB compounds, Life Chemicals kinase inhibitor library compounds, compounds derived through pharmacophore modelling, and Enamine library compounds, respectively.

As can be seen for CTCB compounds in Supplementary Table 3, out of 24 estimates made for ligand-ATP interaction, 18 indicate a negative binding energy, while 19 estimates show a negative ligand-apoprotein binding energy. The prevalence of ligand–ATP interactions across this compound set is notable and was not initially expected to occur to the extent that was observed.

Regarding compounds derived through docking with Life Chemicals kinase inhibitor library, it can be seen that only compounds coming from the docking run that used crystal 6QU3 structure have negative ligand-protein interaction energy, these being 8542 and 13907 (Supplementary Table 4).

As can be seen for the compounds selected through the pharmacophore modelling, they do not exhibit the same extent of ligand-ATP interaction as seen in CTCB compounds, although the ligand-apoprotein binding energy is something that was clearly selected for (Supplementary Table 5). These results are very much in line with the properties that were chosen for in the compounds that contributed to the pharmacophore.

Compounds derived from Enamine library virtual screen are quite interesting in their diversity of performance (Supplementary Table 6). Here, for example, is compound clus6-1, which has the most positive binding energies out of all considered compounds but still managed to stay in the binding pocket for 500 ns of MD simulation without escaping it. We can also some extent of similarity shared in compounds that were classified into the same clusters. For example, the only two compounds showing any evidence of interaction with ATP are found in cluster number 7.

### Correlation of MM-PBSA results and wet-lab derived IC50 values

We have noticed that there is a relationship between our MM-PBSA values and previously calculated IC50 values [6]. Namely, there is a relationship between difference in between ligand-apoprotein and ligand-ATP binding energies (binding energy_ligand-apoprotein_ - binding energy_ligand-ATP_, which we will henceforth call Delta for purposes of simplicity) and IC50 values. The results for 100 ns and 500 ns can be found in Figs 1 and 2. Using these, we have found that we might be able to estimate IC50 values with some accuracy, especially using estimates spanning time from 300 ns to 500 ns from the beginning of simulations. The estimates based on these regressions can be found in Table 2. It is very important to stress the R^2^ values for both regressions, with R^2^ = 0.6047 and R^2^_500 ns_ = 0.8081. The second value, especially, is surprisingly high with possible utility not only in predicting IC50 values but also possibly for generating insights into the mechanism of inhibition.

**Figure 1:**
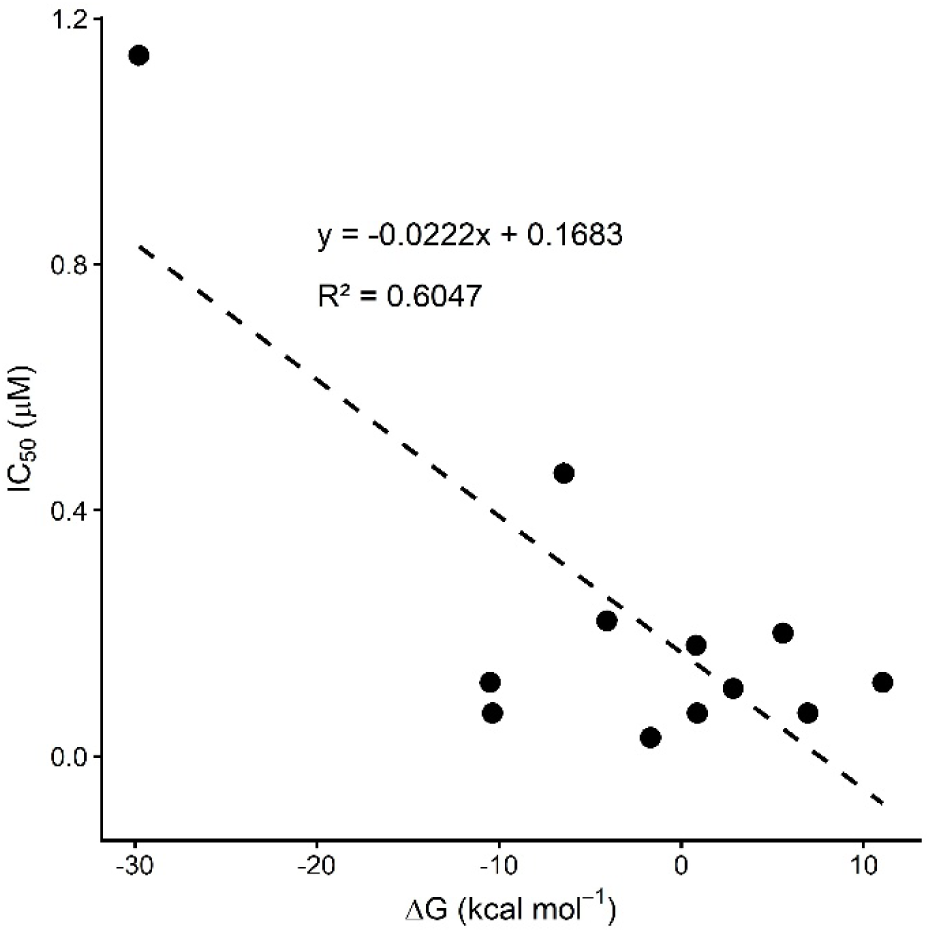
Delta vs IC50 values of CTCB compounds for 100 ns.

**Figure 2:**
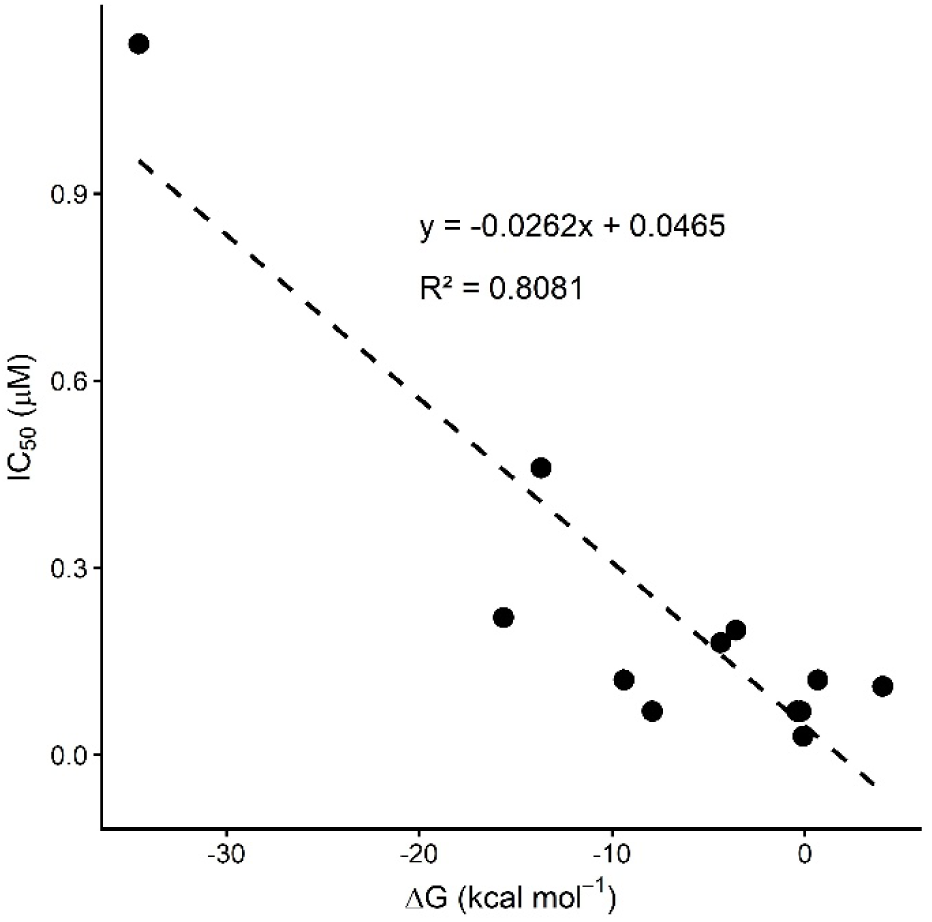
Delta vs IC50 values of CTCB compounds for 500 ns.

**Table 2:**
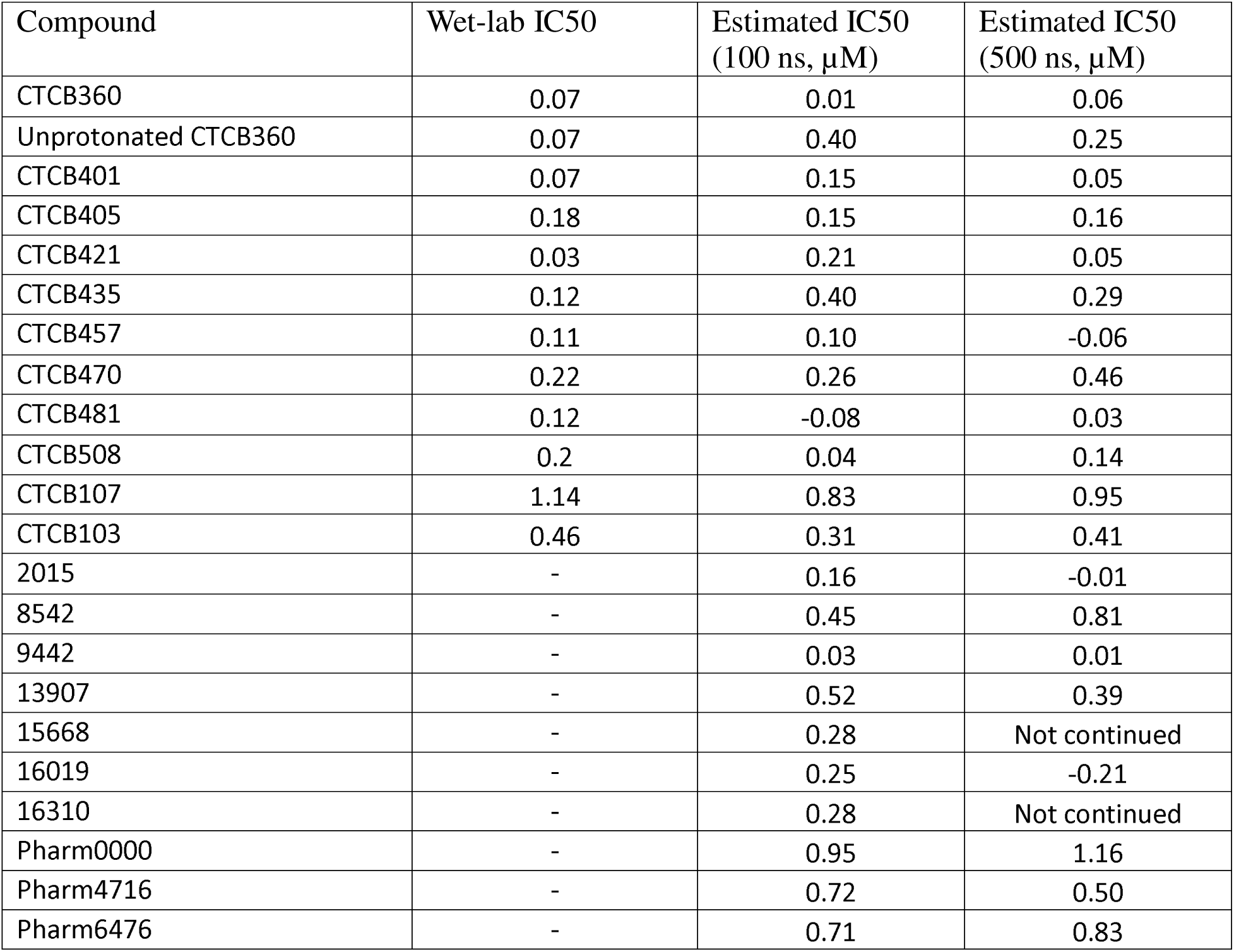
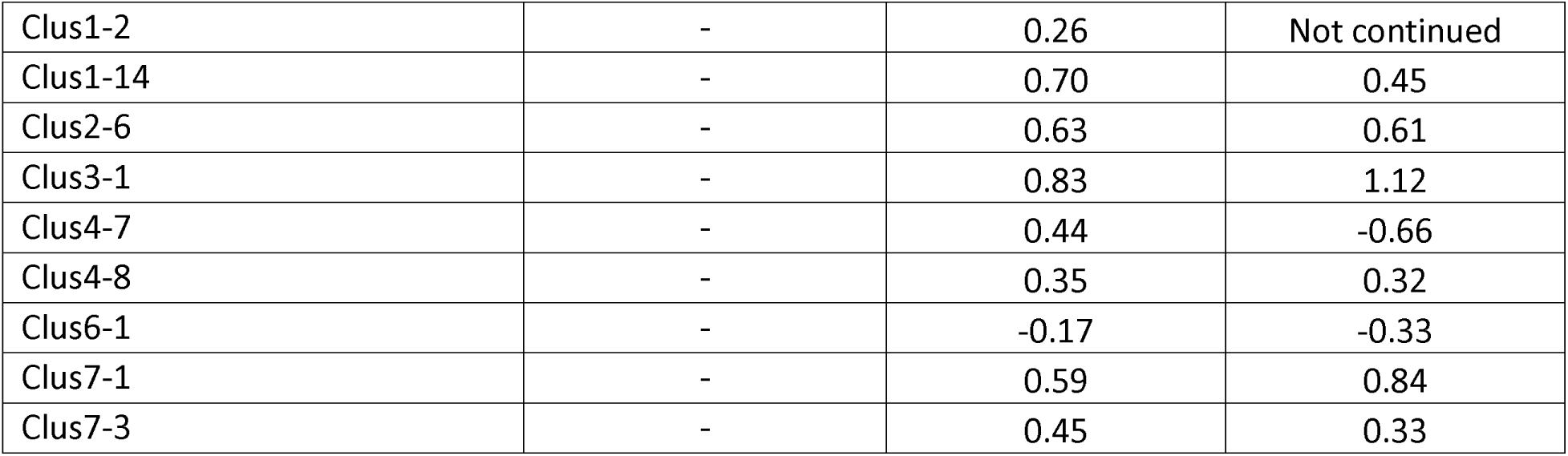
Wet-lab and estimated IC50 values.

### Hydrogen bond analysis

We then analysed hydrogen bonds in the MD trajectories. The results for different compound sets are reported in Supplementary Tables Supplementary Table 7, Supplementary Table 8, Supplementary Table 9, and Supplementary Table 10.

We can see in Supplementary Table 7 that all CTCB compounds form some H-bonds with apoprotein, while some also form H-bonds with the ATP molecule. Specifically, 16 out of 36 estimates show some H-bond formation between the ligand and the ATP molecules.

In the case of Life Chemicals kinase inhibitor library compounds, only two out of nine compounds show some ligand-ATP H-bond formation (Supplementary Table 8). Interestingly, no such interaction for any of the compounds is detected at 500 ns, although the significance of this is unclear.

Similarly to Life Chemicals compounds, only one compound derived from pharmacophore modelling is predicted to form H-bonds with ATP, and this interaction is again lost as simulation progresses beyond 100 ns (Supplementary Table 9). Another interesting and significant finding is that these compounds form more H-bonds than CTCB compounds, which is in line with binding energy differences between these two groups of compounds.

We have observed a relationship between the binding energy and H-bond estimates. We report a relationship between number of H-bonds involved in ligand-apoprotein interaction as well as ligand-apoprotein binding energy (Figures **Figure 3 and Figure 4**). As seen by R^2^ values, the extent to which these are related does not change as simulations progress.

**Figure 3:**
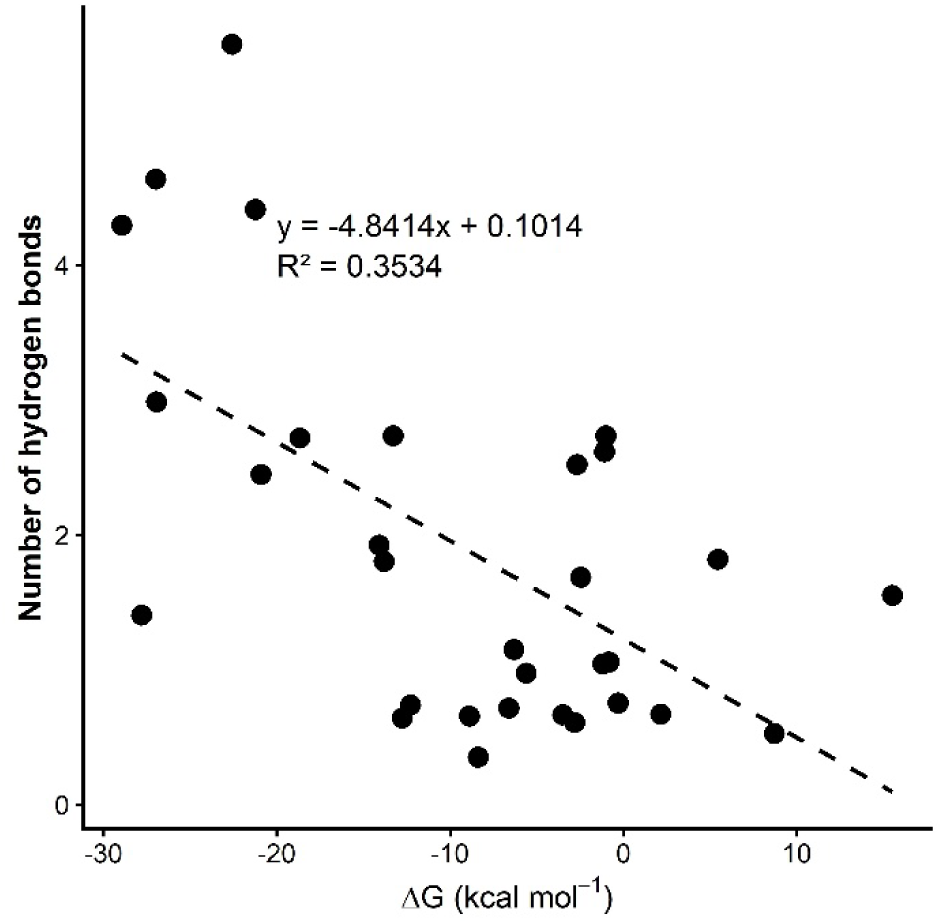
Relationship between predicted number of H-bonds and the predicted binding affinity at 100 ns.

**Figure 4:**
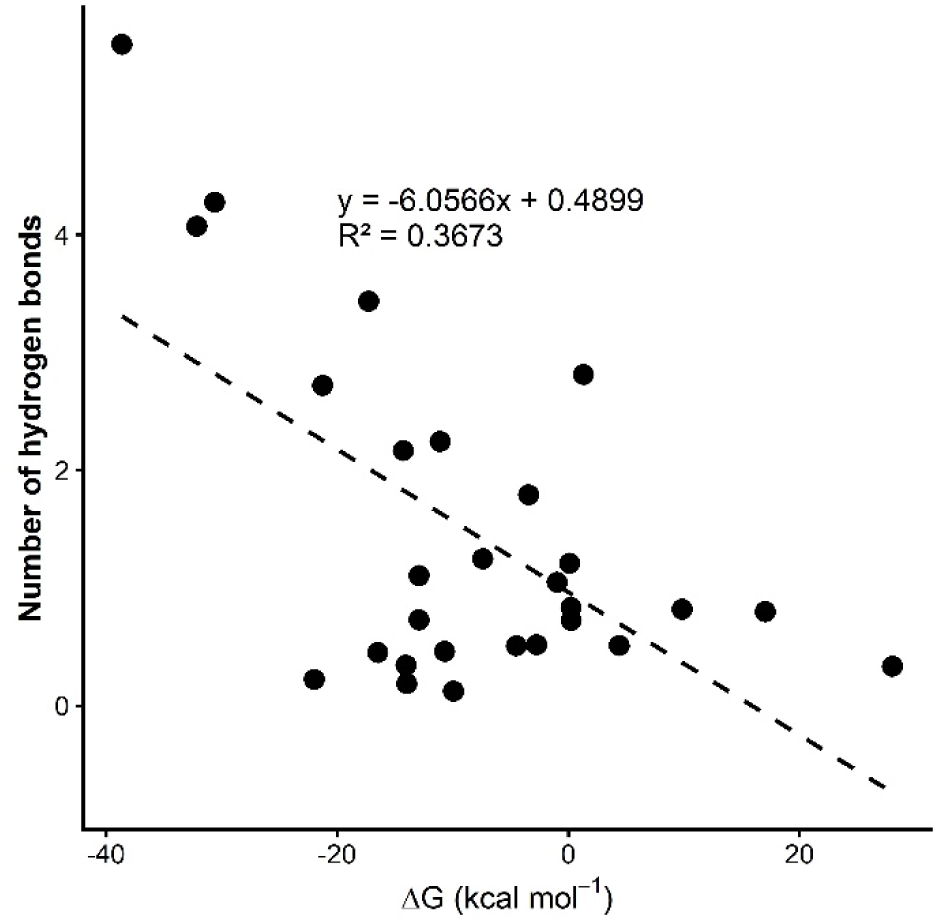
Relationship between predicted number of H-bonds and the predicted binding affinity at 500 ns.

### Prediction of charge at pH 7.4 and isoelectric point

Since pKa values of ligands were reported to have a significant effect on EC50 values, we also decided to analyse these for the compounds that we studied. We have used Chemicalize to estimate the isoelectric point and charge of molecular species at pH 7.4. The results are summarised in .

Table 3. Here N/A means that the software was not able to make a prediction except for 15668 and phram4716, for which the software did not find any ionisable atoms were found. If groups are compared by their average charges, average charge at pH=7.4 of CTCB compounds is 0.62, while those of Life Chemicals kinase inhibitor library compounds, pharmacophore-modelling derived compounds, and Enamine library compounds are 0.29, -0.50, and 0.00 respectively. Therefore, CTCB compounds as a group have the highest charge. The significance of this will be touched upon in the discussion.

**Table 3:**
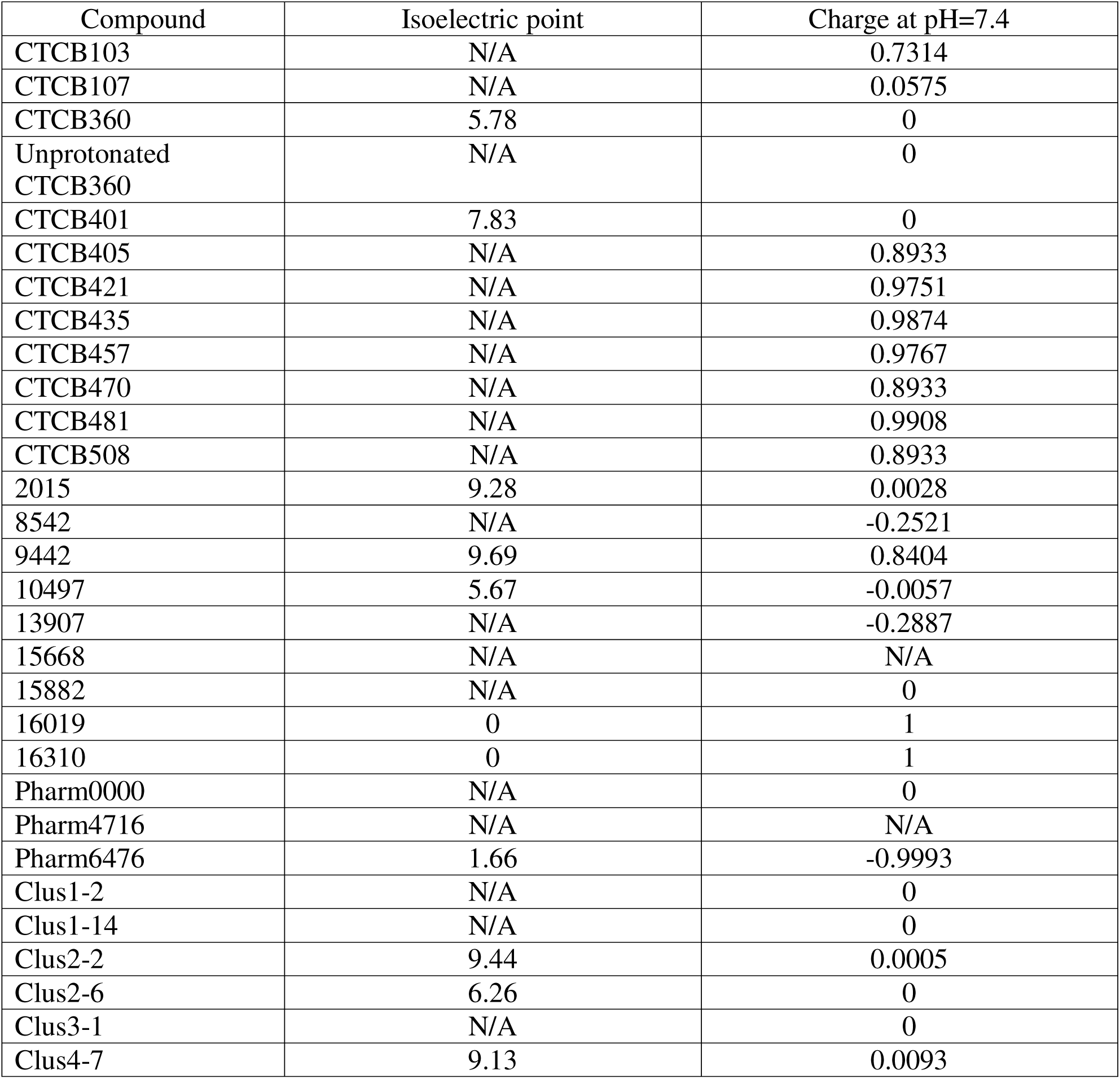
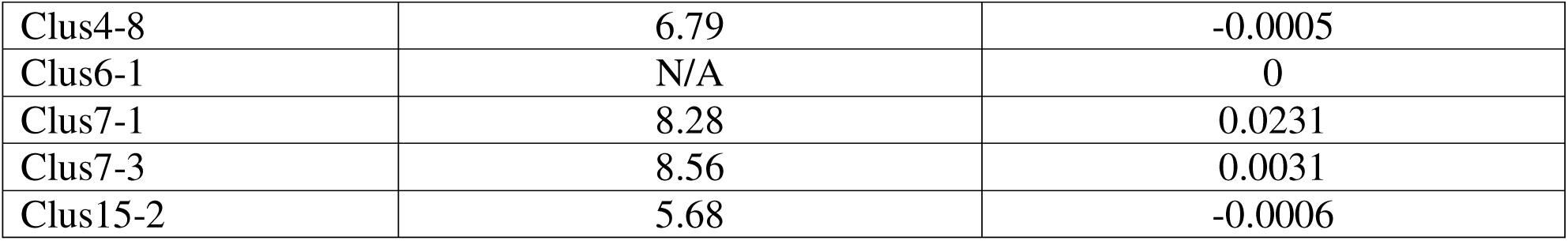
Prediction of the isoelectric point and charge at pH=7.4.

### Selection of the most promising compounds

We used MM-PBSA results, the IC50 estimates based on MM-PBSA results and scoring of Hydrogen bonds to select a subset of ligands that seem the most promising. The results are reported in Table 4. We have assigned points to compounds based on their performance and, looking at the CTCB compound data, chose an arbitrary threshold of 7. The compounds could get one point for negative binding energy in ligand-apoprotein and ligand-apoprotein-ATP complex estimates, two points for IC50 estimated to be not more than 0.25 µM at 100 ns, three points for binding energies in ligand-apoprotein and ligand-apoprotein-ATP complex not more positive than -5 kcal/mol, four points for IC50 estimated to be not more than 0.25 µM at 500 ns and up to 3 points for Hydrogen bonds. In the case of hydrogen bonds, averages for CTCB compounds were made across six estimates, namely ligand-apoprotein and ligand-ATP interactions at 100 ns, 300 ns and 500 ns. Compounds would get 0.3 points for exceeding these averages for estimates at 100 ns and 0.6 points for exceeding these averages at 300 ns and 500 ns. A threshold of 7 points was chosen for inclusion of compounds for further consideration as promising ones, 10 compounds were thus selected: CTCB360, unprotonated CTCB360, CTCB401, CTCB405, CTCB421, CTCB457, 2015, 9442, 16019, and clus6-1.

**Table 4:**
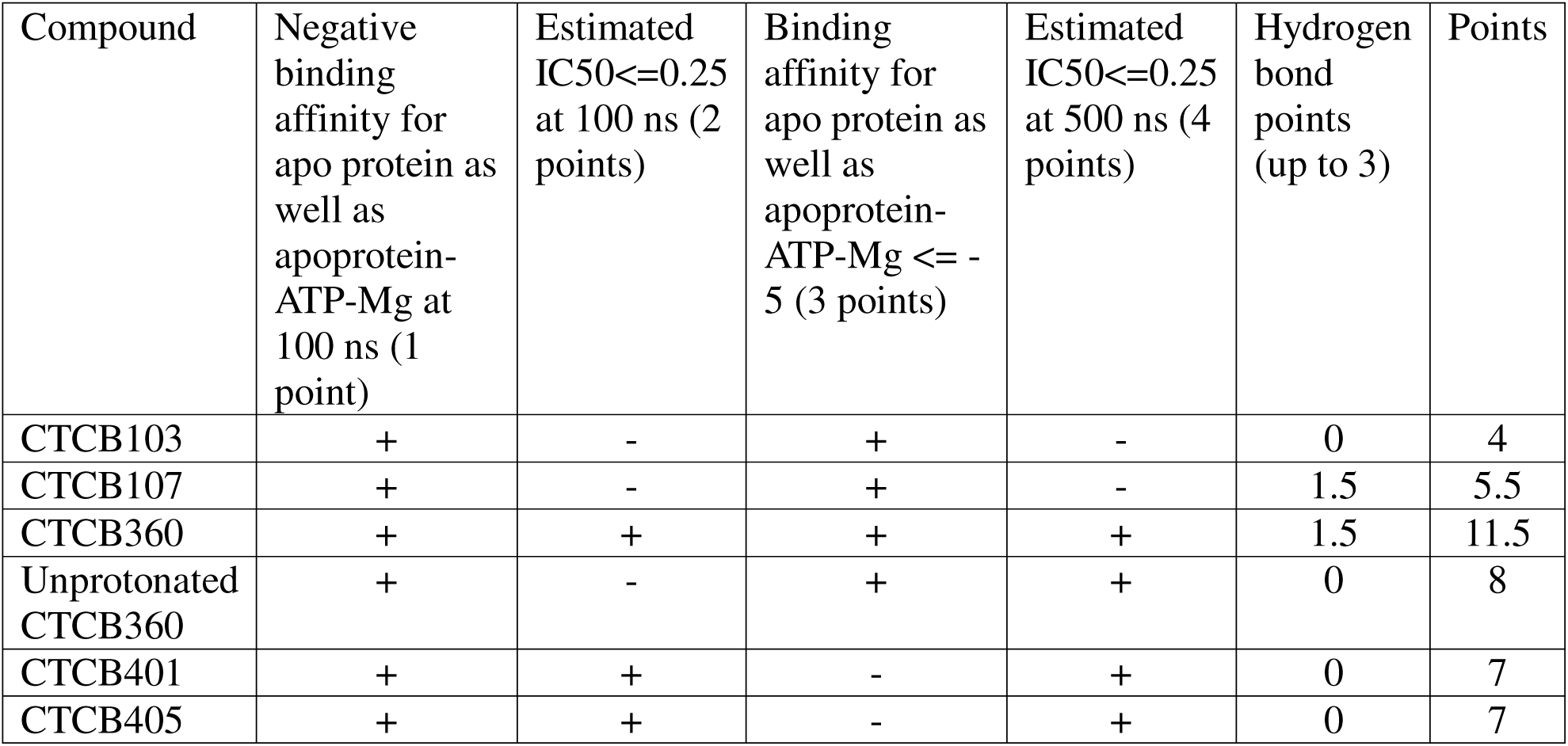
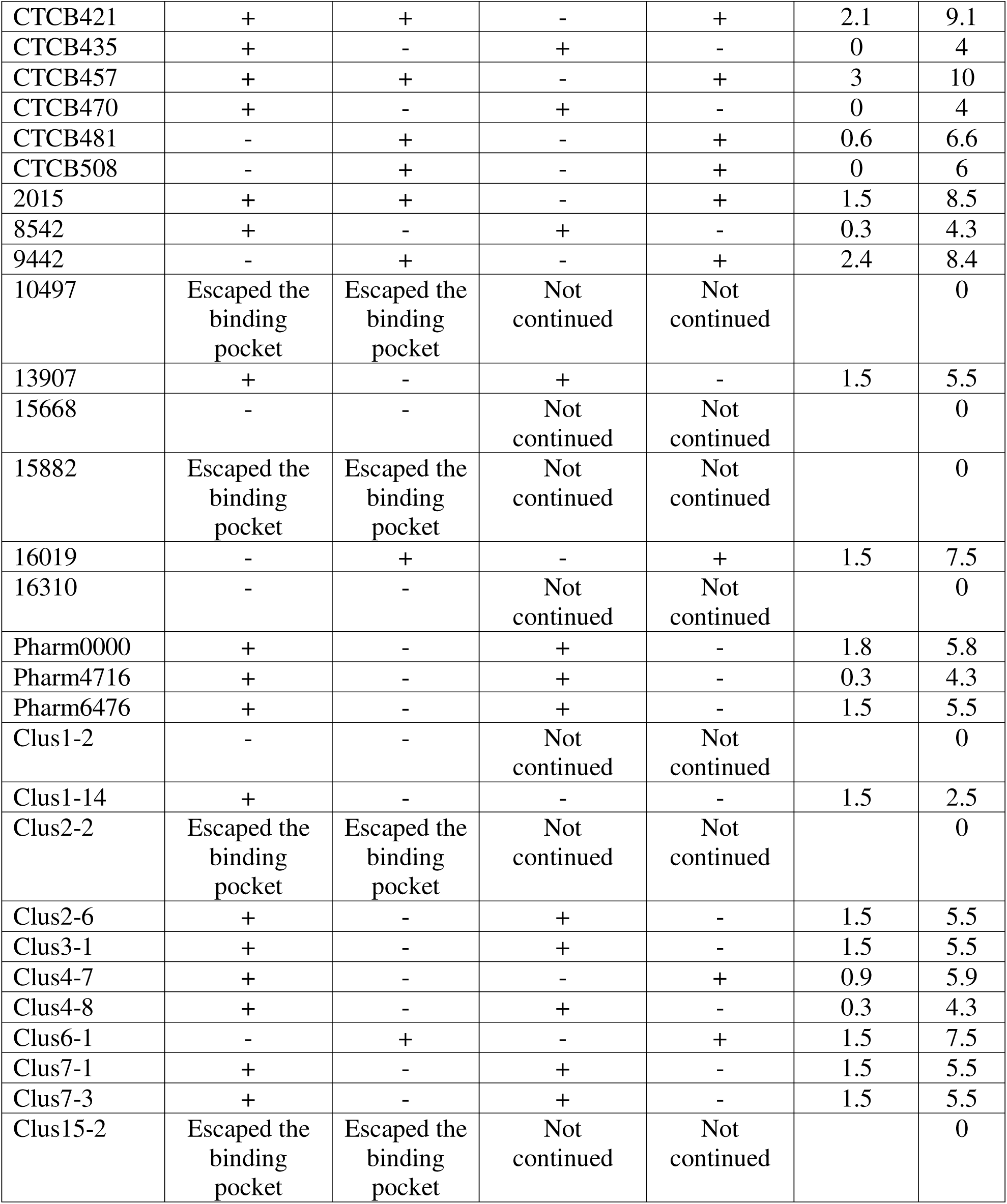
Selection of the compounds.

### ADME prediction

We have used SwissADME and Chemicalize for predictions of drug-likeness and other medicinal chemistry parameters. Namely, MW, TPSA, H-bond acceptor, donor, and rotatable bond counts were estimated using Chemicalize, while everything else was estimated using SwissADME. The results are summarised in Supplementary Table 11 for CTCB compounds and Supplementary Table 12 for non-CTCB compounds.

### Correlation of IC50 values, charges at pH=7.4 and EC50 values and prediction of EC50 values

We have also used linear regression to analyse the relationship between the Delta at 500 ns values, the charge at pH=7.4 and the EC50 values. As mentioned previously, it was already noted by previous studies that the pKa values have a significant effect on the EC50 values [6]. Correlation between predicted and real EC50 values of CTCB compounds is reported in Figure 5.

**Figure 5:**
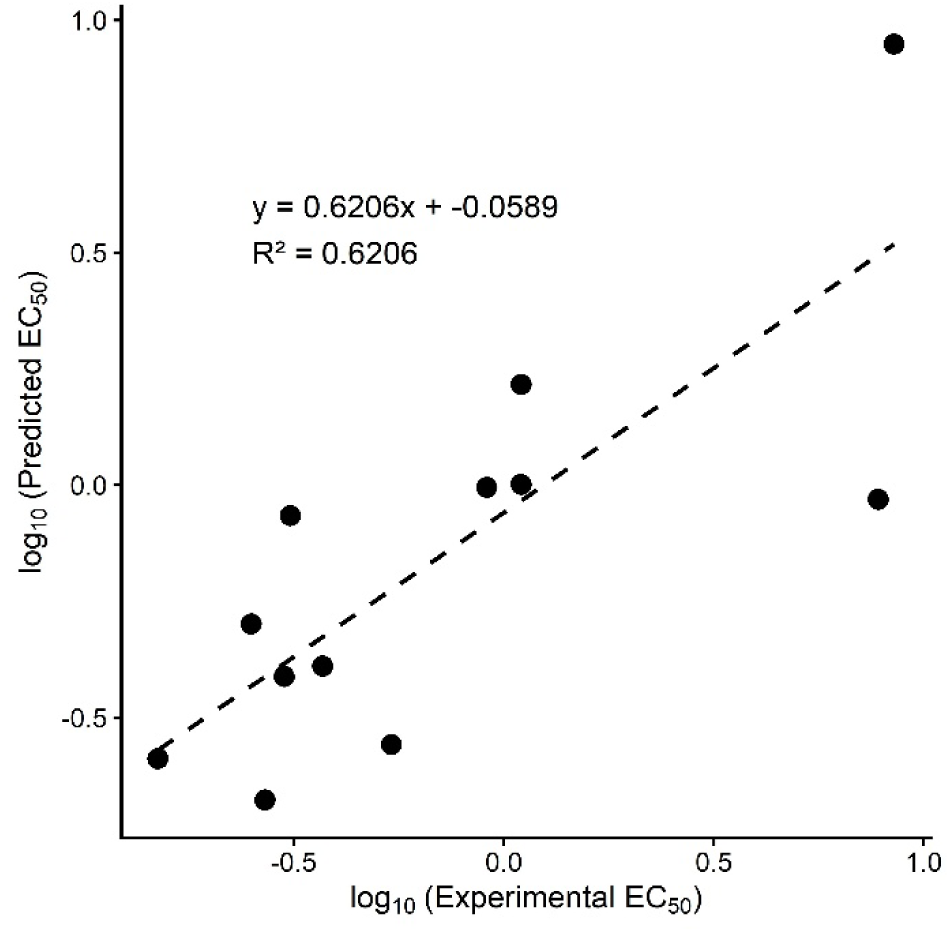
Relationship between predicted and real EC50 values of CTCB compounds.

We did not include these EC50 predictions in the overall appraisal of the compounds and selection of the most promising ones due to the fact that here we have two independent variables and so the regression model might be suffering from overfitting given a relatively small number of data points. However, beyond that, the results of our analysis agrees with the observations of other authors.

We have also used the regression model to predict EC50 values of non-CTCB compounds. The predicted EC50 values are reported in Table 5.

**Table 5:**
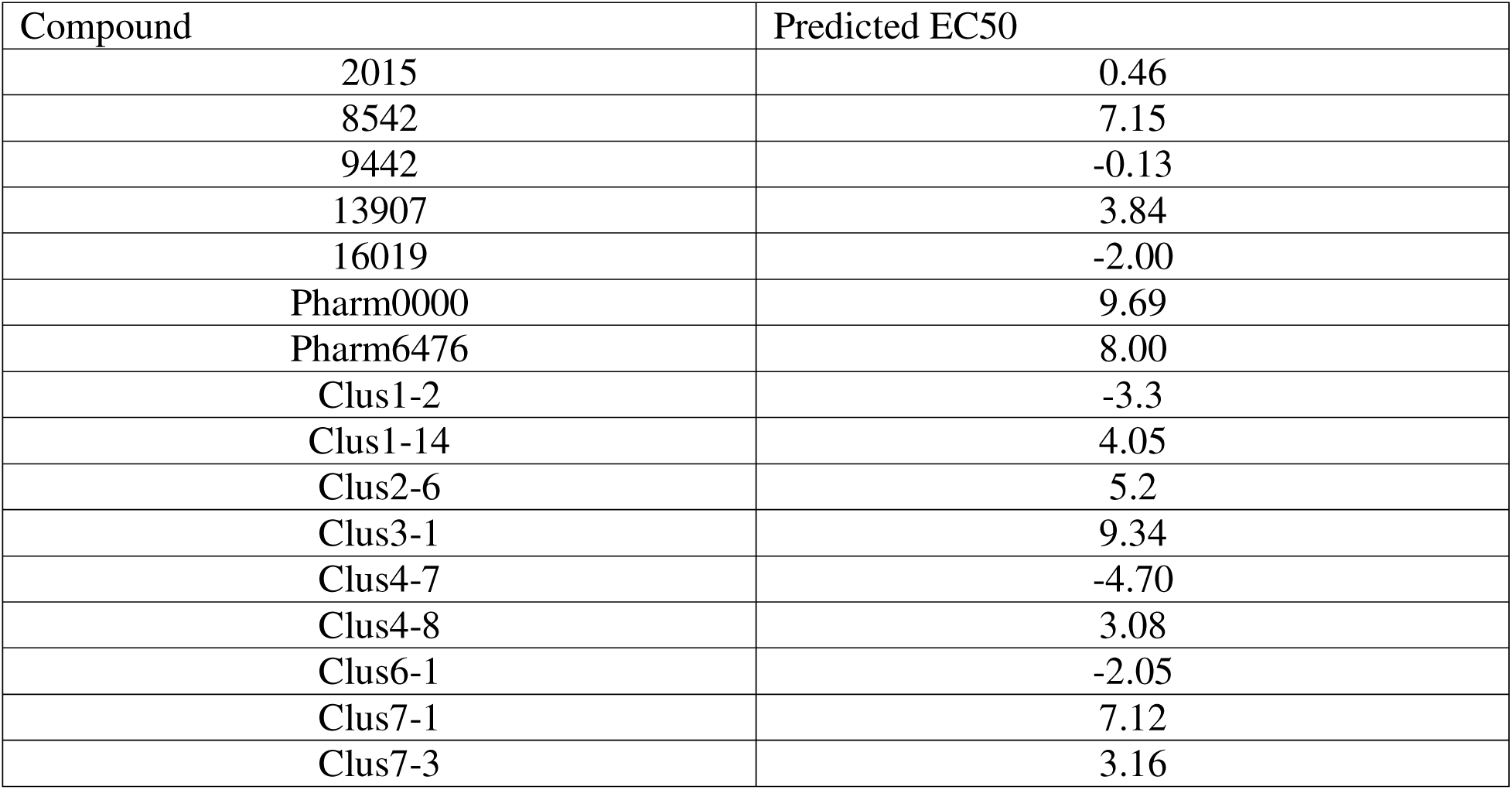
Predicted EC50 values of non-CTCB compounds.

### Identification of ligand-binding pocket

As a next step, we wanted to check the residues that ligands form Hydrogen bonds with. We performed this analysis only for the selected compounds. The results can be found in Supplementary Table 13. We have then tried to define a consensus binding pocket that would be correlated with predicted efficacy of ligands. We have defined this pocket as residues found at least twice in the CTCB compounds as well as at least once in non-CTCB compounds. The thus-defined consensus binding pocket therefore involves the following residues: 15, 17-18, 20, 198-199, 202-203, 226, 231-233, 274-275, 341, 430-434. Afterwards, we used this binding pocket definition to see whether the compounds that we selected led to significantly different behaviour of protein than compounds that were not selected. For that, we calculated and report RMSF data for all residues, and for the binding pocket residues at 100 ns, 300 ns, and 500 ns. We report results all compounds as well as for apoprotein alone in Supplementary Table 14.

We have not been able to detect any relationships between the RMSF values and the binding energy estimates. Therefore, it does not seem that the binding of efficient inhibitors affects the movements of residues that interact with the protein in an immediately apparent way.

Another aspect in RMSF analysis that we wanted to check was whether the residues that were missing from the crystal protein structures that we had to model in behave differently from other residues. Results are reported in Table 6.

**Table 6:**
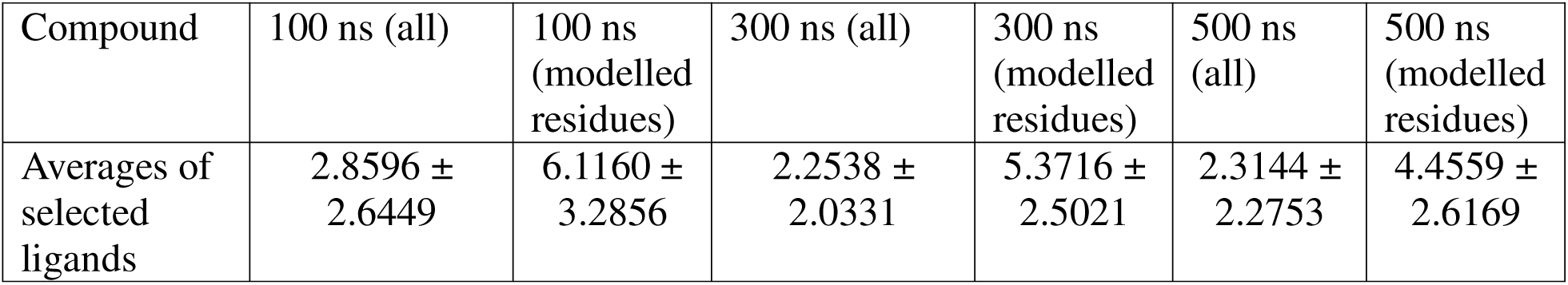
RMSF values for averages of systems containing selected ligands and reported for all residues as well as modelled residues.

### Compound ranking

Since throughout this project we have evaluated many properties of the compounds that we studied, we judged that it would be beneficial to try to rank the selected compounds in terms of performance across all the different metrics that were evaluated. Here, two different ranks are needed, depending on which stage or stages of HAT we are trying to target. Therefore, two rankings, one for stage 1 HAT, and another for stage 1 and 2 HAT are reported.

In case of ranking for stage 1 HAT, blood-brain barrier permeability is a somewhat undesirable trait which could increase the likelihood of side-effects. For stage 1 HAT ranking, following system was devised: points from Table 4 were taken, rescaled to 30 points according to the maximum and regarded as evaluation of ligand-enzyme interaction, EC50 values were also considered, additional points were granted for solubility, lack of CYP inhibition, drug-likeness, and lack of blood-brain barrier crossing. Compounds could acquire up to 100 points. Results are reported in Table 7.

**Table 7:**
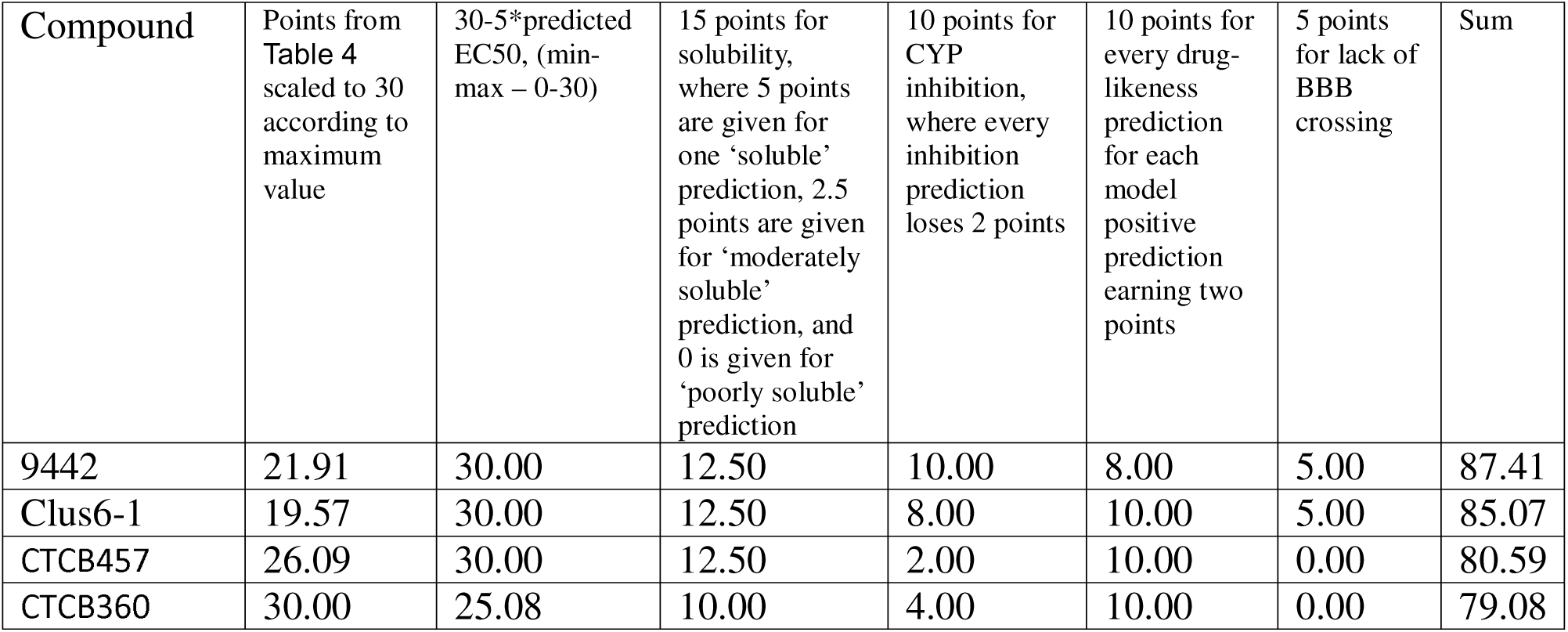
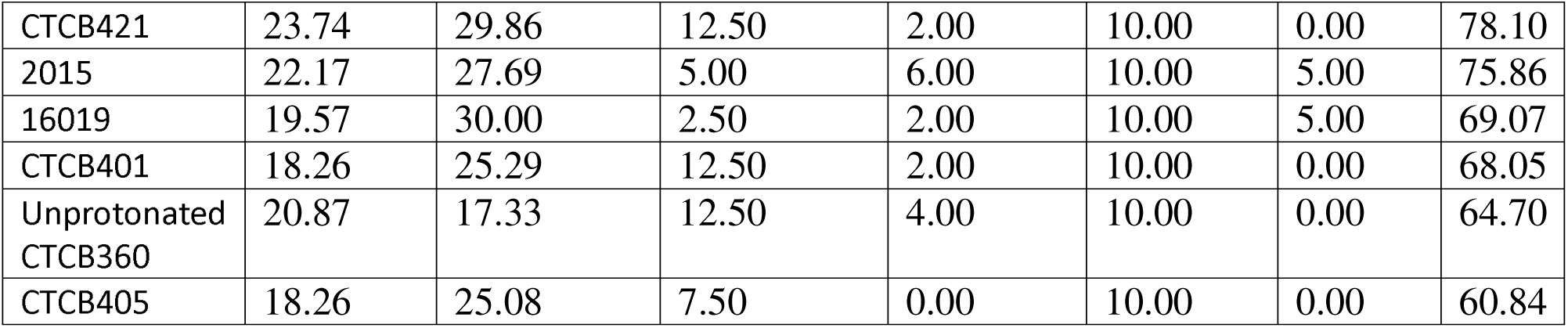
Ranking of compounds for stage 1 HAT.

In contrast, in the case of ranking for both stage 1 and stage 2 HAT, blood-brain barrier permeability is a key characteristic. Therefore, a different ranking system was developed: points from Table 4 were taken, rescaled to 20 points according to the maximum, EC50 values were also considered, additional points were granted for solubility, lack of CYP inhibition, drug-likeness, and blood-brain barrier crossing. Compounds again could be scored up to 100 points. Results are reported in Table 8.

**Table 8:**
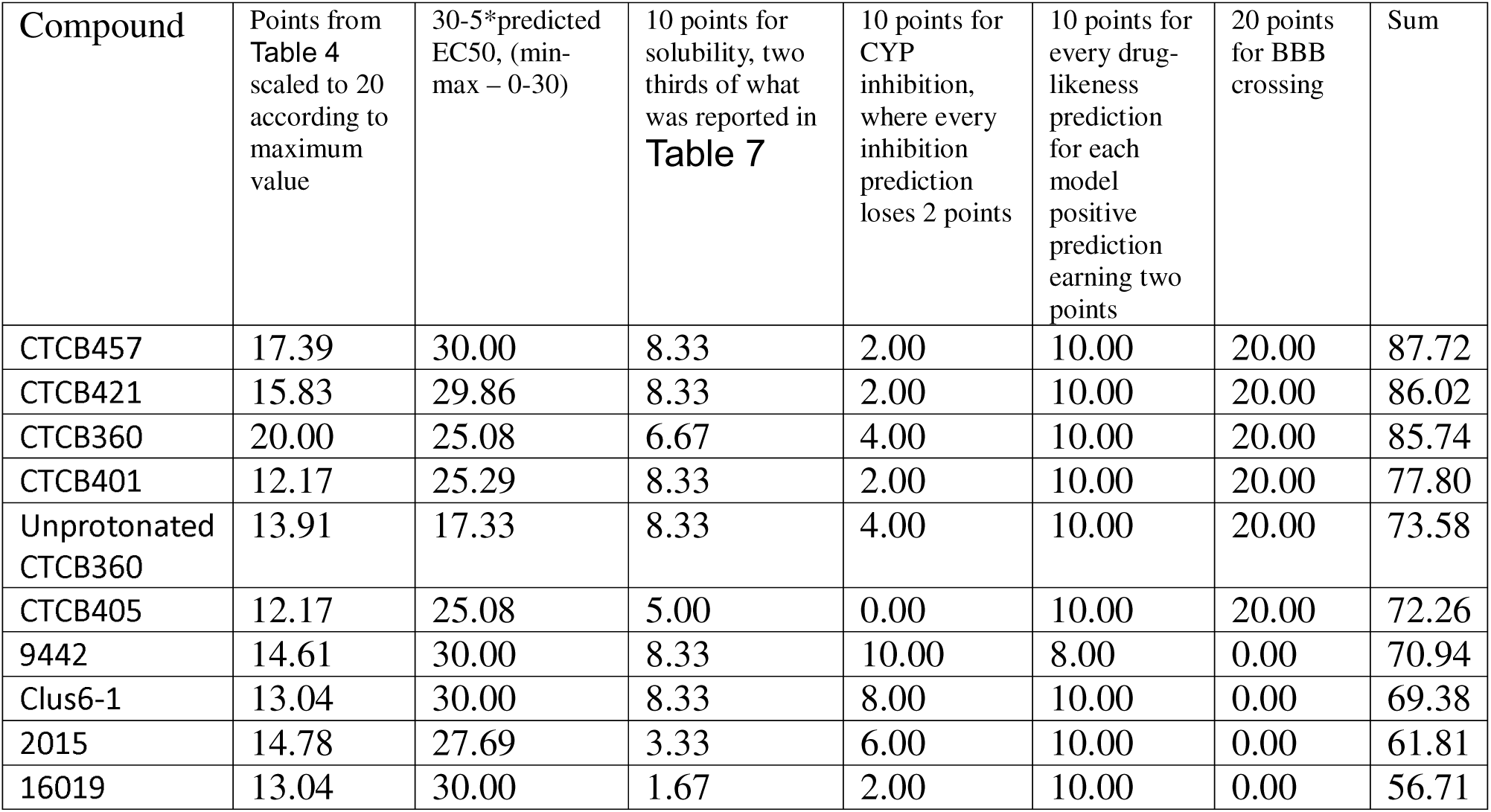
Ranking of compounds for both stage 1 and 2 HAT.

Interestingly, according to the ranking system that we devised here, for stage 1 HAT, several compounds, namely 9442 and clus6-1 outperform all CTCB compounds. 2015 and 16019 rank in the middle of all compounds. The best-performing CTCB compounds in the stage 1 HAT setting are CTCB457, CTCB360, CTCB421.

Ranking compounds for both stage 1 and stage 2 HAT, CTCB compounds have a clear advantage over non-CTCB compounds due to ability to cross the blood-brain barrier. This is not surprising, the lack of this ability in non-CTCB compounds leaves them unable to reach *T. brucei* residing in CNS.

## DISCUSSION

Since this project analysed various pools of compounds and used different sets of methods for different compounds, we made several charts to summarise the workflow. Figure 6 shows the workflow used to produce the pharmacophore and Figure 7 shows the workflow that was used on all the sources of compounds to evaluate them.

**Figure 6:**
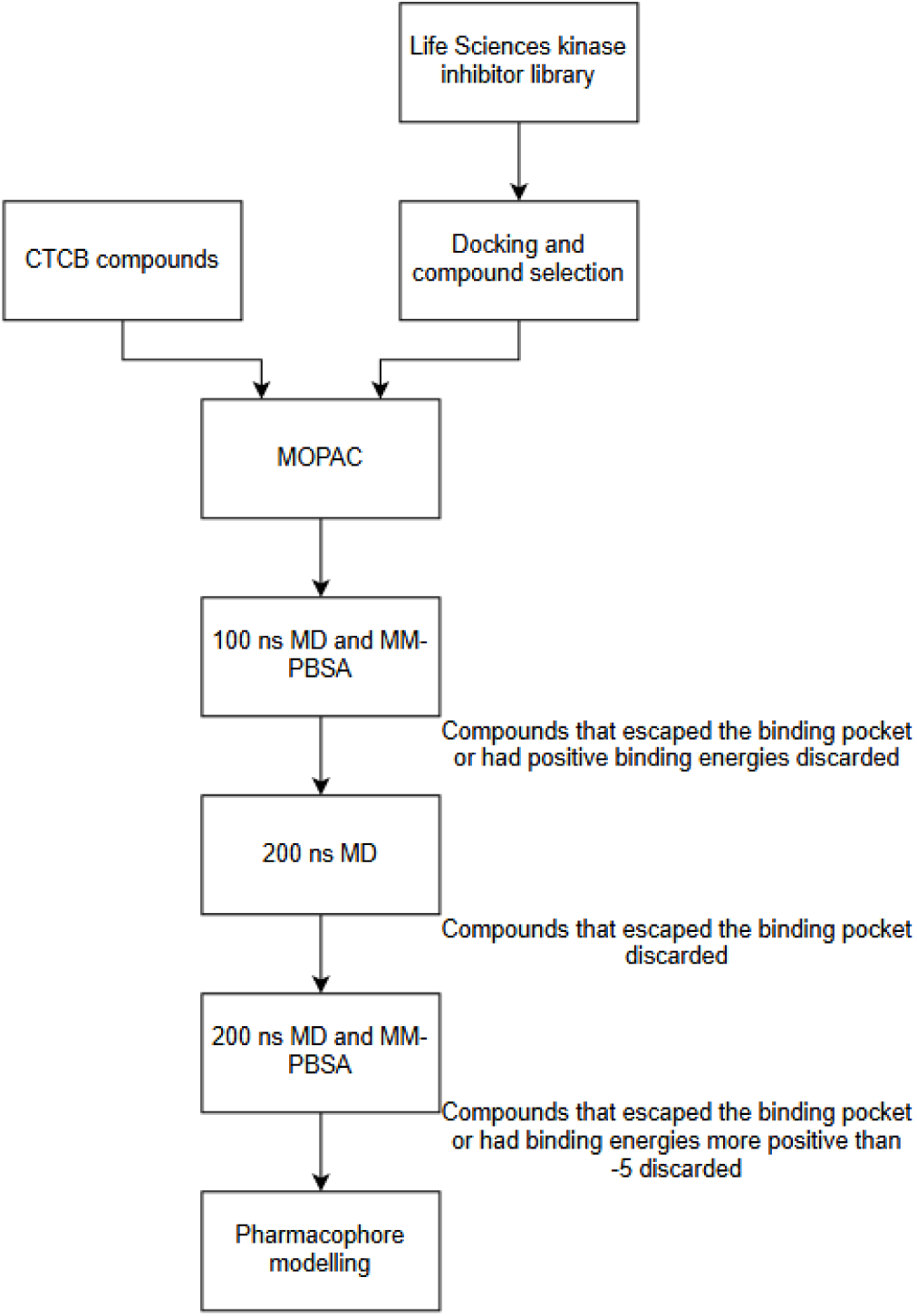
Steps used to produce the pharmacophore.

**Figure 7:**
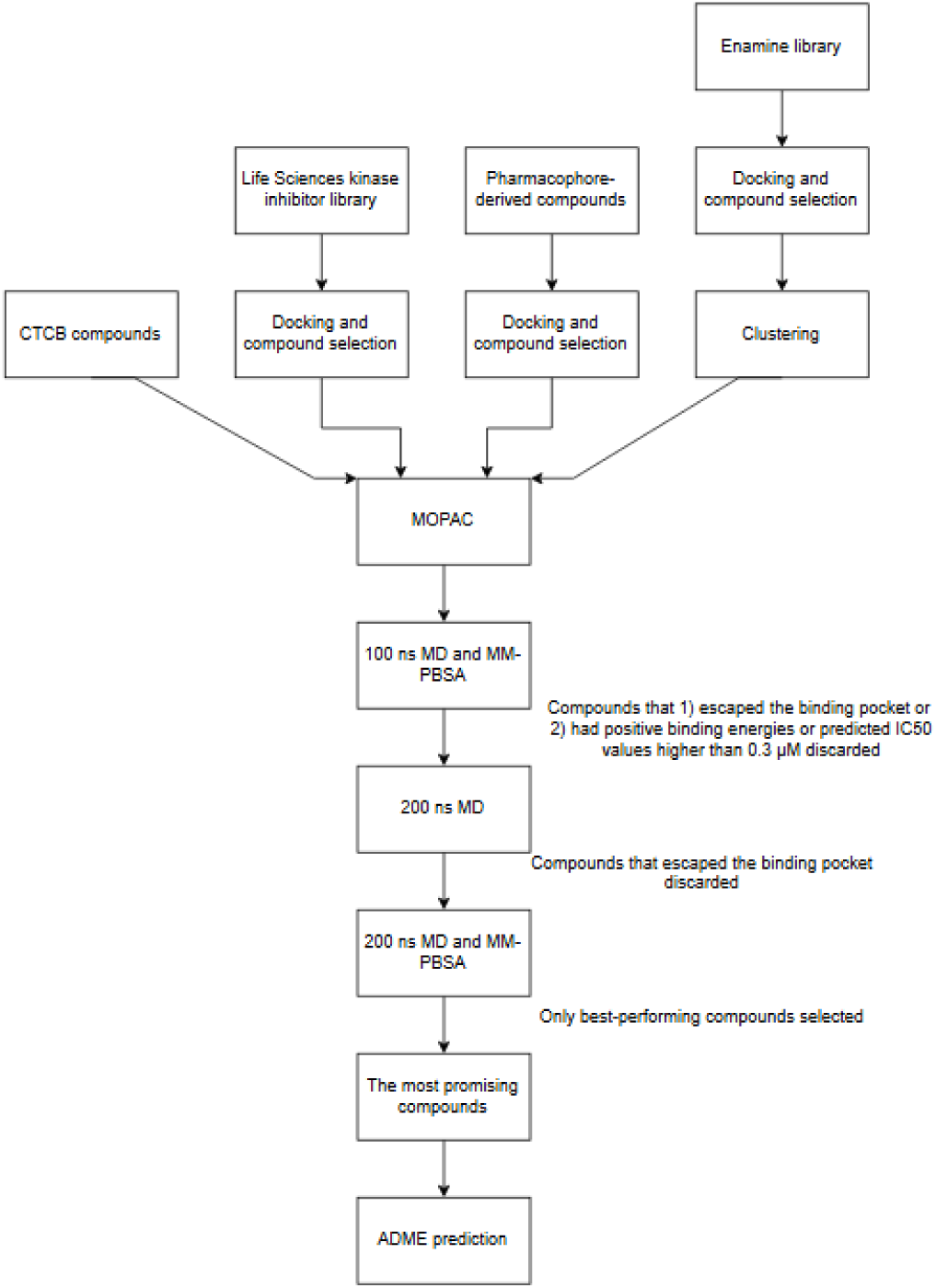
Steps followed to select the most promising compounds.

Interestingly, in our project, we found that ATP played an important role in the overall performance of the small molecule inhibitors. Specifically, we have found that the difference between the binding energy of the small molecule and the apo protein and the small molecule and the ATP molecule is highly predictive of the IC50 values. To emphasise, more negative inhibitor-ATP binding energies and more positive inhibitor-apoprotein binding energies lead to higher inhibitory effects. Several aspects of this are noteworthy. First and foremost, the degree to which this difference correlates with IC50 is remarkable and unexpected. In light of this, our simulations appear to be highly realistic and potentially possess substantial predictive power when used on novel molecules. Secondly, this is unusual in the sense that when designing small molecule inhibitors, researchers usually favour molecules that tightly bind to the protein in some key site. Our data would suggest that our system is an example in which this sort of thinking would not lead to optimal results. Thirdly, this insight specifically in the case of TbPFK is a novel one, to the best of our knowledge, previous studies have not analysed the role of inhibitor-ATP interaction in TbPFK inhibition.

To expand on the accuracy of the simulation, there are several aspects that should be mentioned that make the results of simulations convincing. First, the already mentioned correlation between the binding energies and the IC50 values is crucial. Secondly, the direction and extent to which Hydrogen bonds correlate with binding energies is also convincing, that is, we would expect higher hydrogen bond counts to lead to more negative binding energies and for these two to correlate somewhat moderately, as there are also other kinds of interactions contributing to the binding energies, our data is in line with these expectations. Thirdly, the residues that were not present in the crystal structures that we have used and that we had to model in as we made our systems are also the residues that during the simulations tend to move the most. This again agrees well with our expectations, as the protein parts that are flexible would be anticipated to be missing in crystal structures. Utility of MM-PBSA is still contested when compared to other methods, however, it appears to have been accurate in our project [37]. This reasonably satisfactory performance in our case could have been due to 1) quite long simulation times as compared to what many publications pursue, and 2) SEQM use prior to MD runs.

It is also important to emphasize a very significant false positive rate of virtual screen protocols. Previous literature highlights that as low as only 12% of top compounds derived through virtual screens show activity in *in vitro* assays [38]. This represents a substantial challenge in employing these methods. In case of our project, several steps were taken to try to mitigate this. First, as much as possible, we tried to combine several pieces of docking software together or, when using only one, combined them with another method, such as molecular clustering. In case of use of several different pieces of docking software, the hope is that if a compound shows good performance in two different tools, the false positive possibility is decreased. Whereas for the use of clustering, the hope here was to try to find molecular patterns that are associated with good performance and trying to further evaluate as diverse patterns as possible. Ultimately, the subsequent use of MD, a method supposed to be more accurate than docking, leads to higher reliability of selected compounds. In our case, we also had CTCB compounds already tested *in vitro* and *in vivo* [6]. These compounds were useful for us to try to evaluate additional compounds screened and allowed us to make predictions of biological activity of the screened compounds.

Previous research noted a relationship between pKa values of compounds and their EC50 values. We ourselves have evaluated this and found, even though our dataset for evaluation might have been small and prone to overfitting, that there is indeed a relationship between charge at pH=7.4 and EC50. The pH value we chose was different from one chosen previously, which was 7.8. Our choice of pH was based on previous research, which lead us to suspect that *T.brucei* glycosome pH could be expected to be around 7.4 during infection [39].

Furthermore, the blood-brain barrier permeability of drugs targeting *T. brucei* specifically is another aspect that requires consideration. As mentioned previously, there are two stages of HAT, the first being haemolymphatic stage and the second being meningoencephalitic stage [2]. In the second stage, there is brain involvement. Therefore, for molecules to have any activity against stage 2 HAT, the molecule has to permeate the blood-brain barrier. CTCB compounds are able to cross this barrier, and they were shown to have a very high effect for stage 1 HAT model and some effectivity but not absolute parasite clearing in the model involving brain infection. So here, we have to discuss that promising non-CTCB compounds would be good only for stage 1 HAT, while CTCB compounds could be used for stages 1 and 2 with lower effects in stage 2.

Importantly, the 4 selected non-CTCB compounds that we judge to be promising for future studies are predicted not to be able to cross blood-brain barrier. This is a notable limitation of these compounds, since, if the predictions hold true in a lab-setting, these compounds could then only be used to treat stage 1 HAT. However, this limitation is not critical, since if these compounds prove to be well-tolerated by patients and as highly effective or nearly as highly effective as our predictions suggest, they could still be used advantageously in clinical practice. Additionally, compounds able to cross the blood-brain barrier, when used in stage 1 disease, might prove to have more side effects than the compounds that do not cross this barrier.

As mentioned earlier, many authors stress the need for drugs against *T. brucei* that could be used perorally [14,15]. High GI absorption is predicted for almost every compound that we selected as highly promising, with the exception of 16019.

Now let’s quickly go through strengths and weaknesses or other noteworthy points regarding each selected compound:

- CTCB360 displays a favourable drug-like and lead-like profile. This compound is able to cross the blood-brain barrier and thus can be used for both stage 1 and stage 2 HAT. One significant issue is inhibition of CYP1A2, CYP2C19, and CYP2C9, which could indicate that the compound is prone to drug-drug interactions.
- Unprotonated CTCB360 is similar to the protonated form in terms of its drug and lead-likeness. Cytochrome P450 inhibition is also a major issue here. The compound is able to cross the blood-brain barrier.
- CTCB401 also displays drug and lead-likeness. This compound is an inhibitor of 4 evaluated CYPs so the drug-drug interaction risk is even more significant. The compound is able to cross the blood-brain barrier.
- CTCB405 is not lead-like, it also displays the poorest solubility out of all CTCB compounds. It is also the compound that inhibits all CYPs for which predictions were made.
- CTCB421 displays drug and lead-likeness. Overall, it is very similar to CTCB401, with the same strengths and weaknesses.
- CTCB457 is similar to CTCB421 but it is not lead-like.
- 2015 has a fairly high MW of 427.49 g/mol. It is not lead-like. 2015 has a more favourable CYP inhibition profile compared to CTCB compounds. It is unable to cross the blood-brain barrier so it probably can only be used against stage 1 HAT.
- 9442 is not lead-like. There is also one Brenk alert associated with this compound, which could indicate propensity for toxicity, reactivity, and instability, namely, a diketo group [40]. It is predicted to be the only compound with no CYP inhibition.
- 16019 has several unfavourable properties. Firstly, it is predicted to have low GI absorption, making oral administration unsuitable. Secondly, two models predict it to have poor solubility. It also has three Brenk alerts and inhibits four out of five CYPs.
- Clus6-1 is not lead-like. It is only predicted to inhibit CYP3A4.

Another point to discuss is the behaviour of previously defined catalytic residues Asp229 and Asp231. While defining the consensus binding pocket for the selected compounds, based on our inclusion criteria, we included Asp231 in this consensus binding pocket. Previous studies show that T-to-R transition in TbPFK involves these residues, it is also thought that allosteric inhibitors of TbPFK should stabilise inactive conformation of the enzyme. Our inclusion of Asp231 into the consensus binding pocket are compatible with this proposed mechanism of inhibition. As Asp231 interacts with the inhibitors, its certain conformation might indeed get stabilised and prevent this and adjacent residues from taking part in large conformational shifts.

It should be acknowledged, that given the ADME prediction results, 16019 should be discarded from further consideration due to the fact that two different models predicted it to be poorly soluble, even if we were to not pay attention to other potential issues with this compound. Its best use could, perhaps, be as some a starting point which could then be modified to produce better ADME performance, since this compound was predicted to highly effective as TbPFK inhibitor. Given that the target organism at the life stage that we are focusing on is found in the blood, it is crucial for potential drug candidates to exhibit sufficient solubility.

Instead of discarding any more compounds, we instead decided to rank them out based on their overall performance across a range of criteria. Two rankings were made, one for stage 1 HAT and another for both stage 1 and 2 HAT. These rankings revealed some key differences in terms of potential utility of CTCB and non-CTCB compounds. Namely, CTCB compounds have a clear advantage if stage 2 HAT is also aimed to be targeted. In stage 1 HAT context, several non-CTCB compounds are predicted to overperform CTCB compounds, specifically 9442 and clus6-1. Moving forward, wet-lab evaluation of these compounds would be highly useful.

We already touched upon catalytic residues and their involvement in conformational transitions of TbPFK, but it is also important to mention the finding that inhibitor-ATP binding energy seems to be highly correlated with the wet-lab inhibitory activity of the compounds. It is difficult to guess whether our finding and the previous observations are related or if these are different and simultaneous mechanisms by which an inhibitor would affect TbPFK. Whichever is the case, our data clearly indicate that inhibitor-ATP binding is crucial for the inhibitory effects of the compounds. Furthermore, even though we did not observe differences in the apoprotein and ligand-apoprotein complex consensus binding pocket residue RMSF values, the lack of observed difference could be due to insufficiently long MD simulations. Accelerated MD simulations could potentially be used to reveal differences in behaviour.

In our project, through the virtual screens, we have selected the compounds with the lowest inhibitor-apoprotein binding energies. We have also selected compounds with low ligand-apoprotein and ligand-apoprotein-ATP binding energies to build the pharmacophore. These characteristics, as shown by the correlations reported in this paper, might not be best suited for selection of TbPFK inhibitor. Future studies could pursue selection of compounds after virtual screens and for building of pharmacophores using different criteria from what is reported here, for example, lowest estimated IC50 or EC50 values. This perhaps would not return as many compounds that bind with very low binding energies to the apoprotein while possibly not having much inhibitory activity.

## CONCLUSIONS

In this study, we have conducted thorough virtual screening efforts combined with SEQM simulations and MD runs of up to 500 ns. These results have allowed us to make a series of observations about the protein system that we were working with as well as the compounds that we were evaluating. Importantly, evaluating previously wet-lab-characterised compounds has allowed us to attempt to predict behaviour of novel compounds. Beyond having found several novel molecules that outperform CTCB compounds, several additional aspects of our results are particularly interesting: we report very substantial *in silico*-*in vitro* result correlation which validates our approach, we also report an interesting and seemingly key role that ATP plays in the mechanism of allosteric inhibition of TbPFK.

## SUPPLEMENTARY FIGURES

**Supplementary Table 1:**
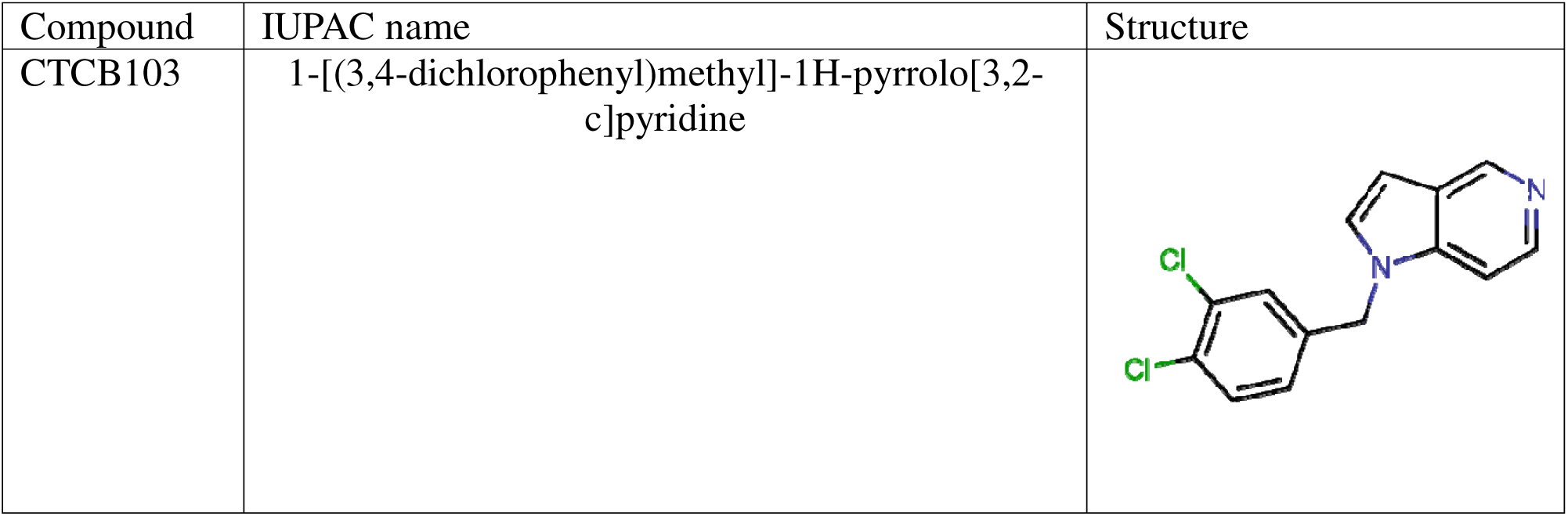
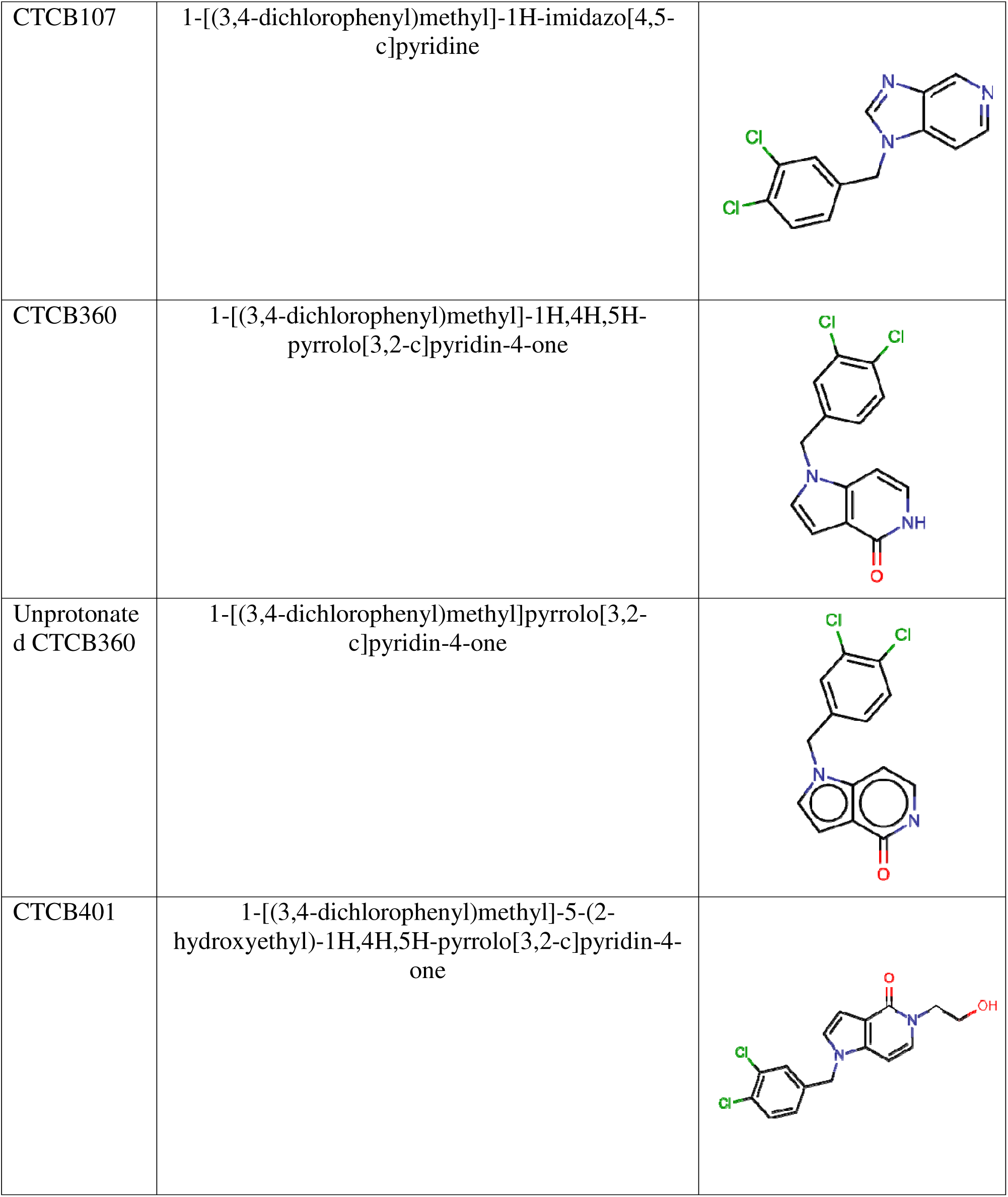
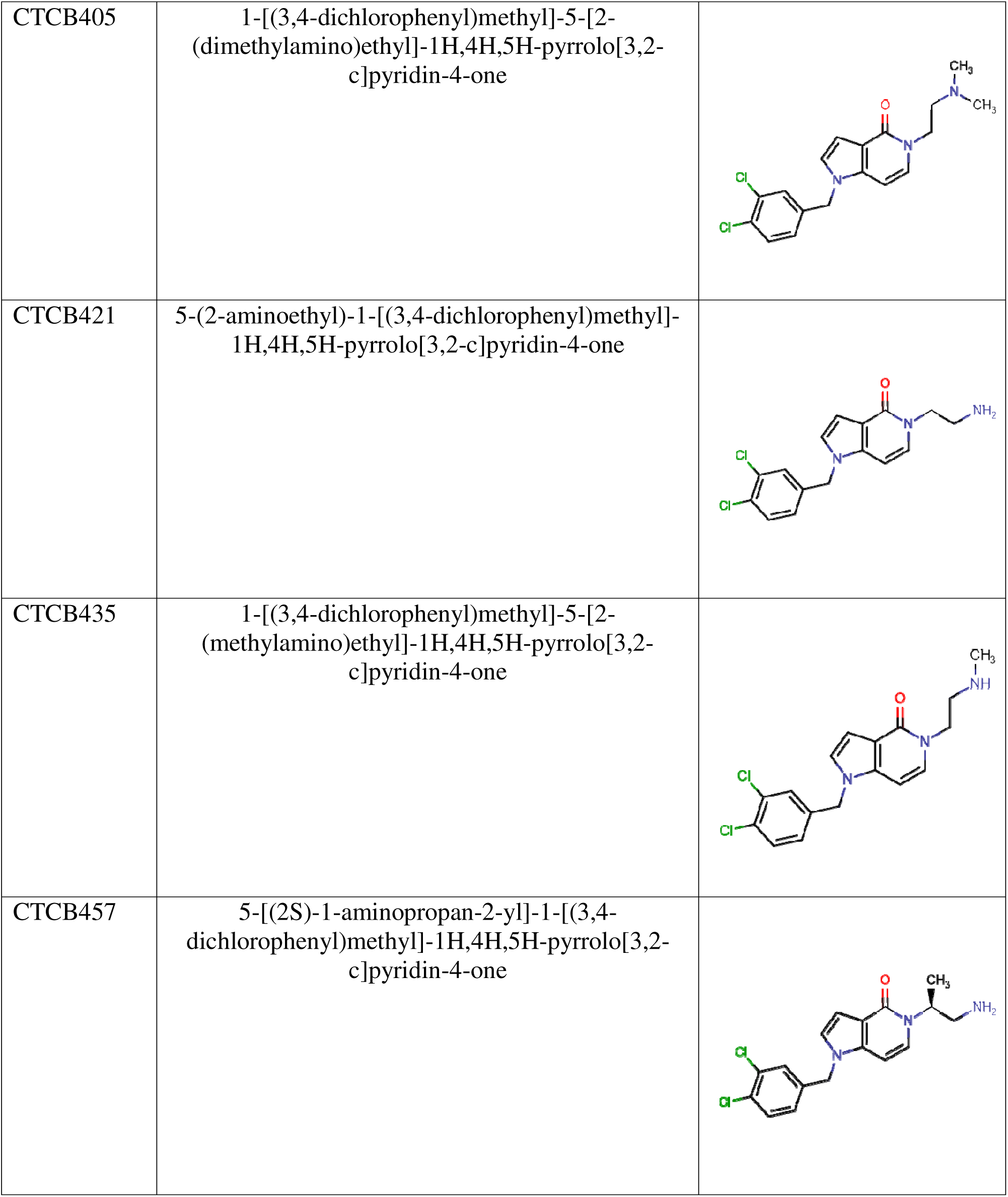
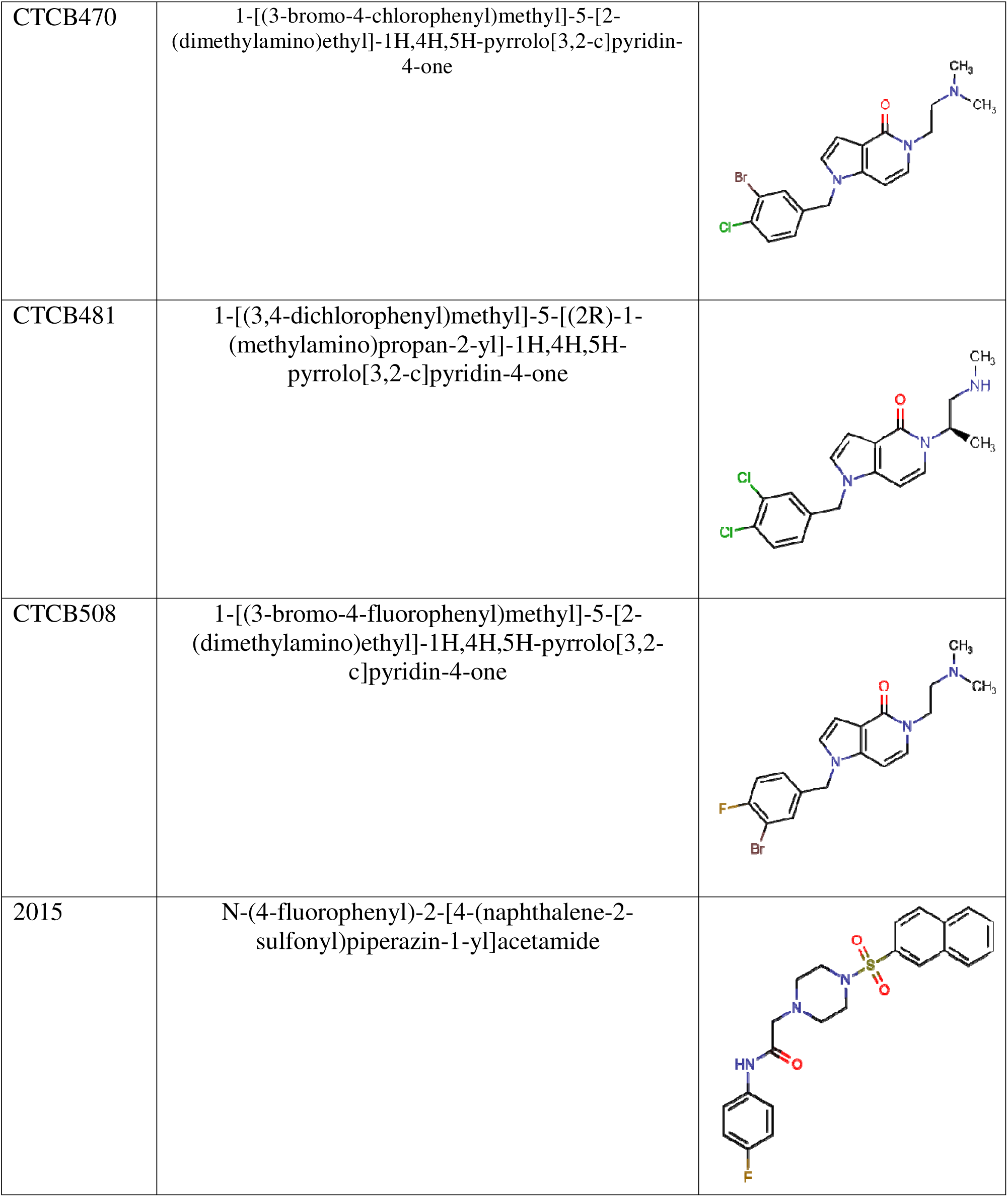
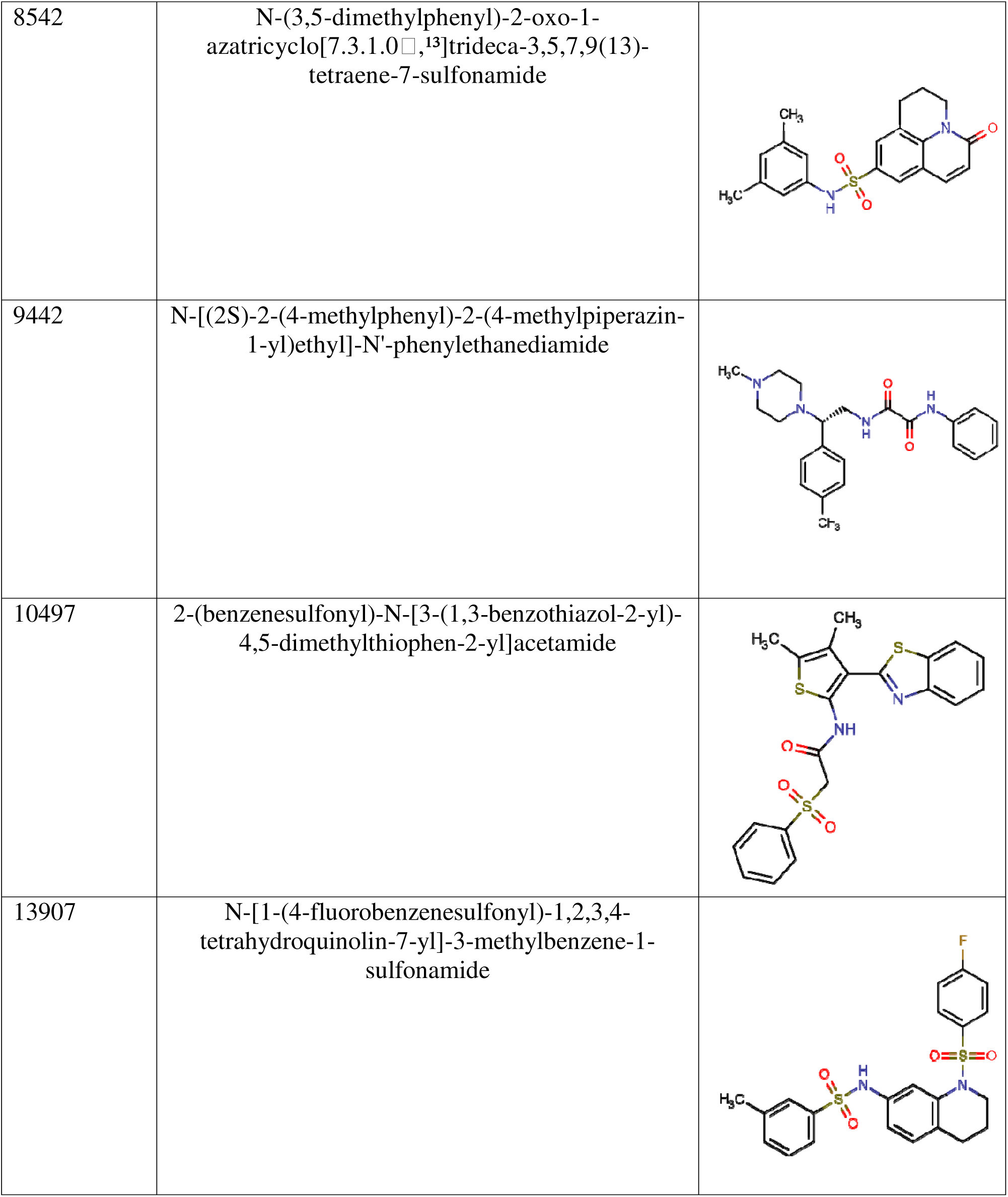
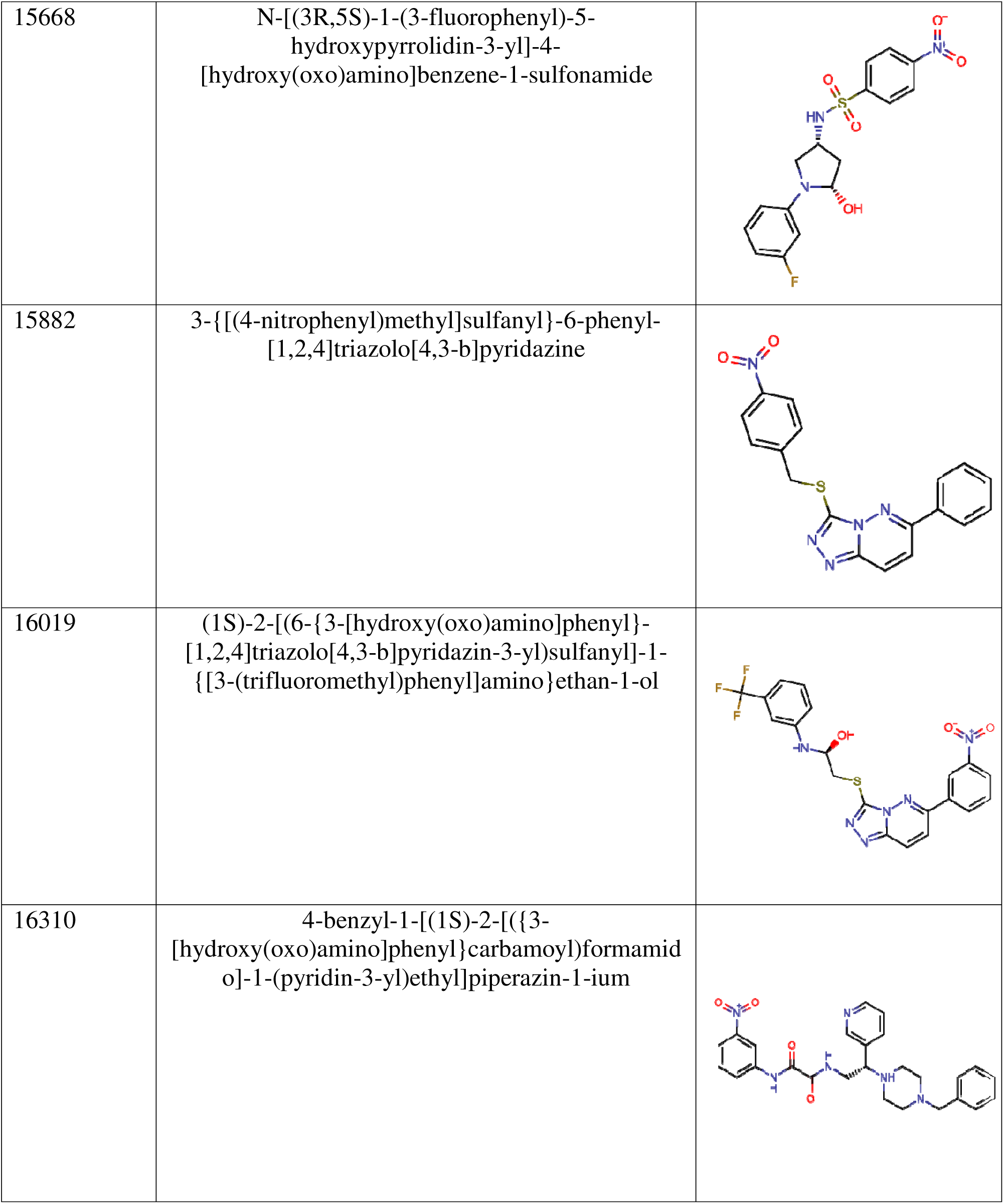
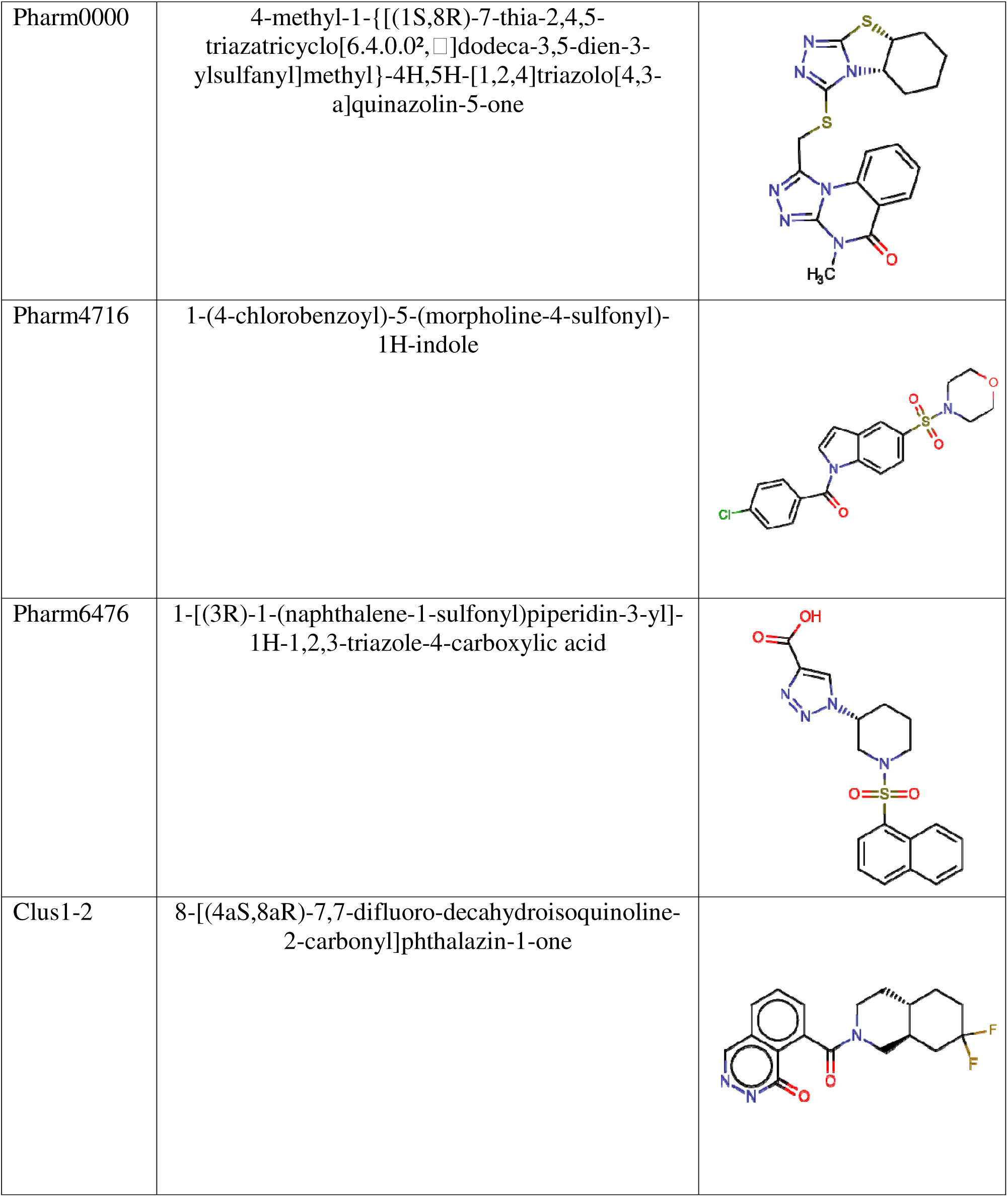
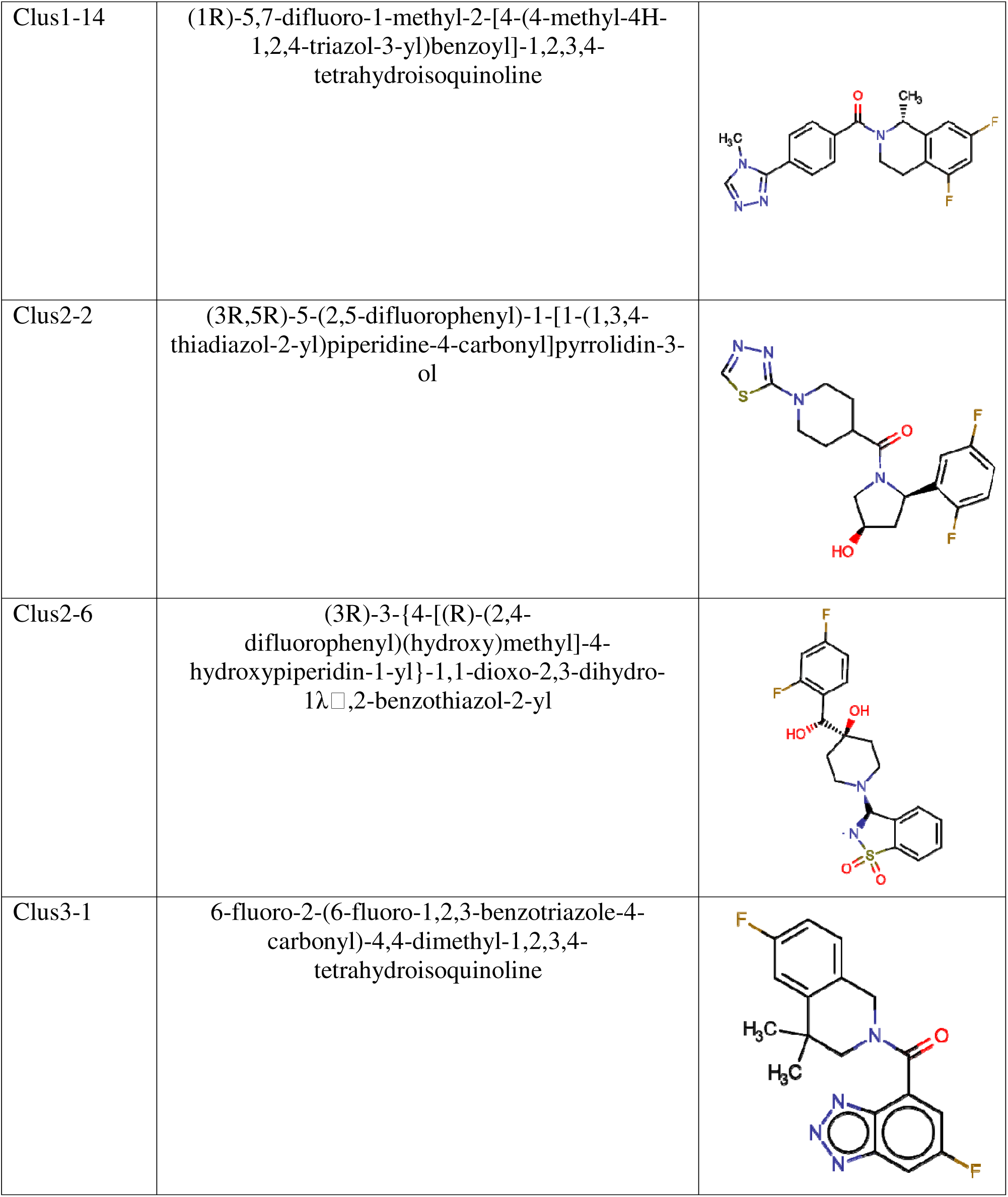
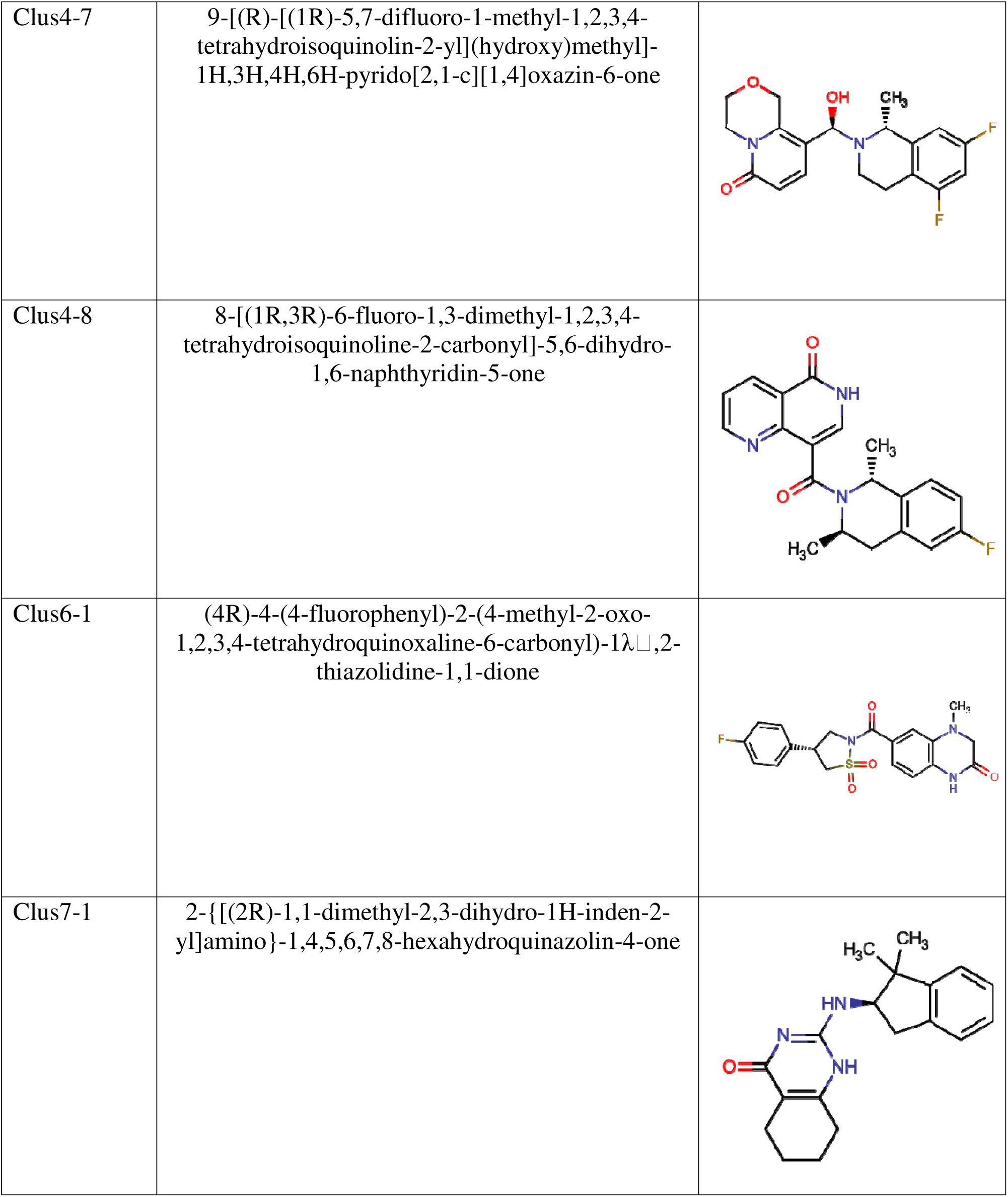
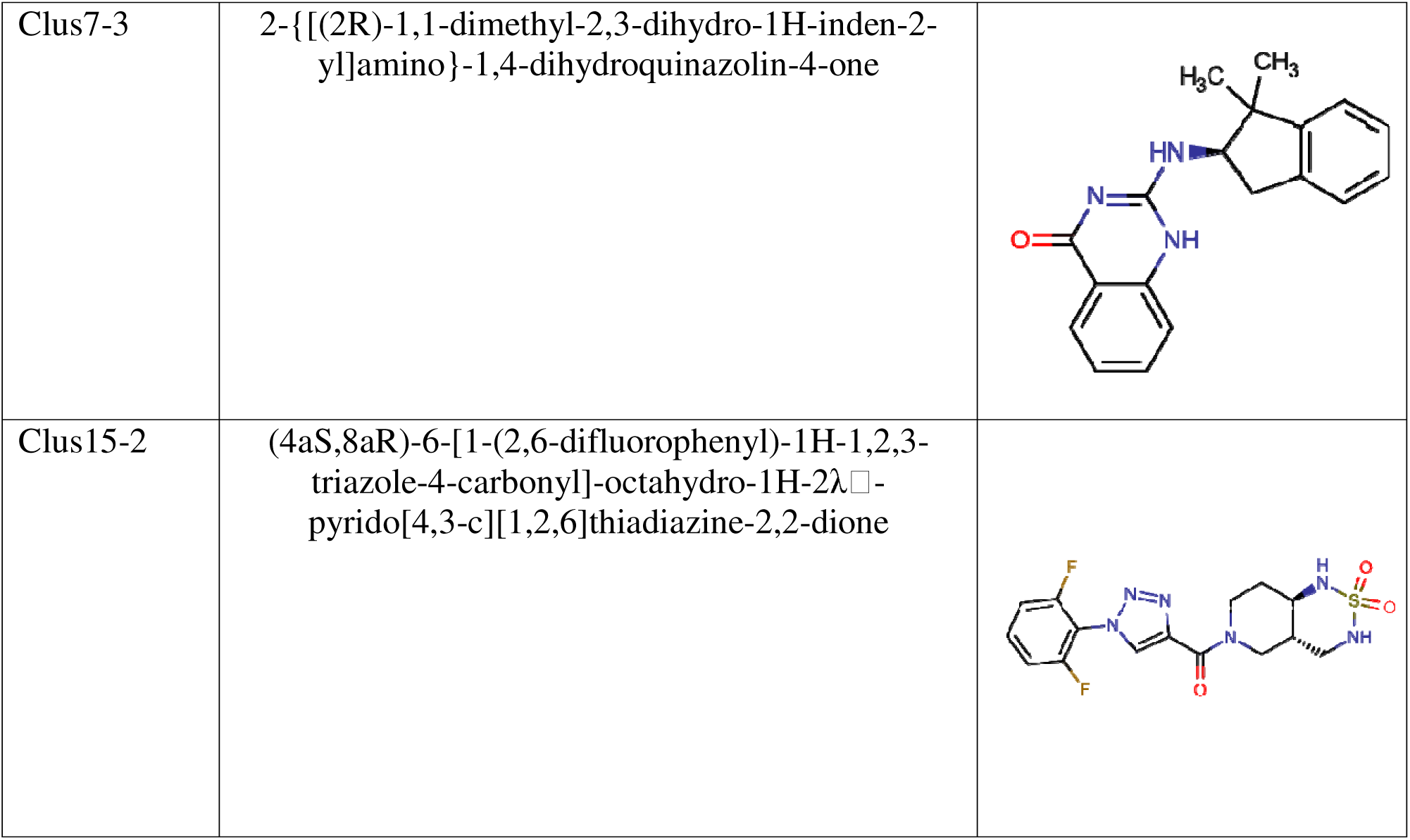
Information on compounds studied.

**Supplementary Table 2:**
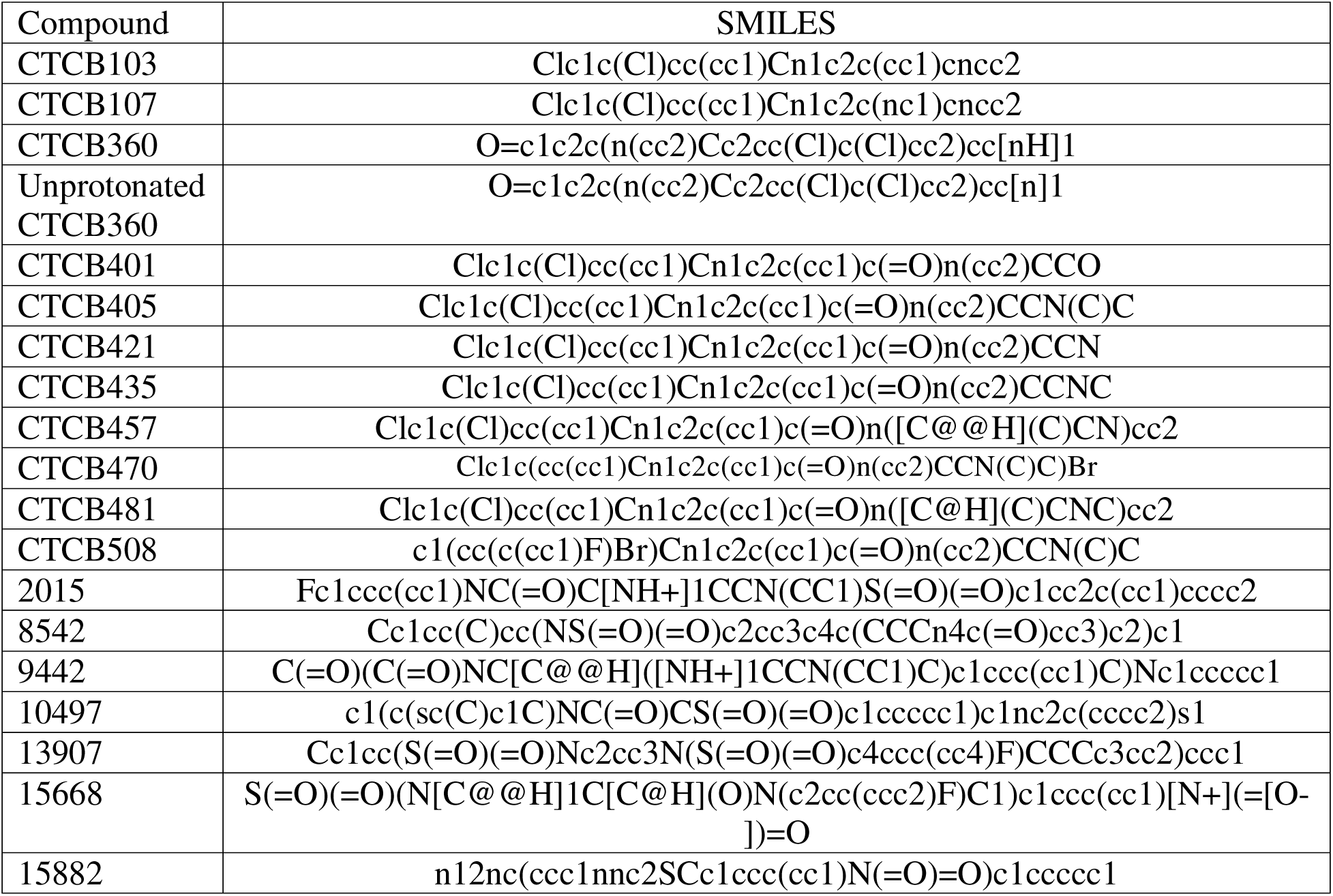
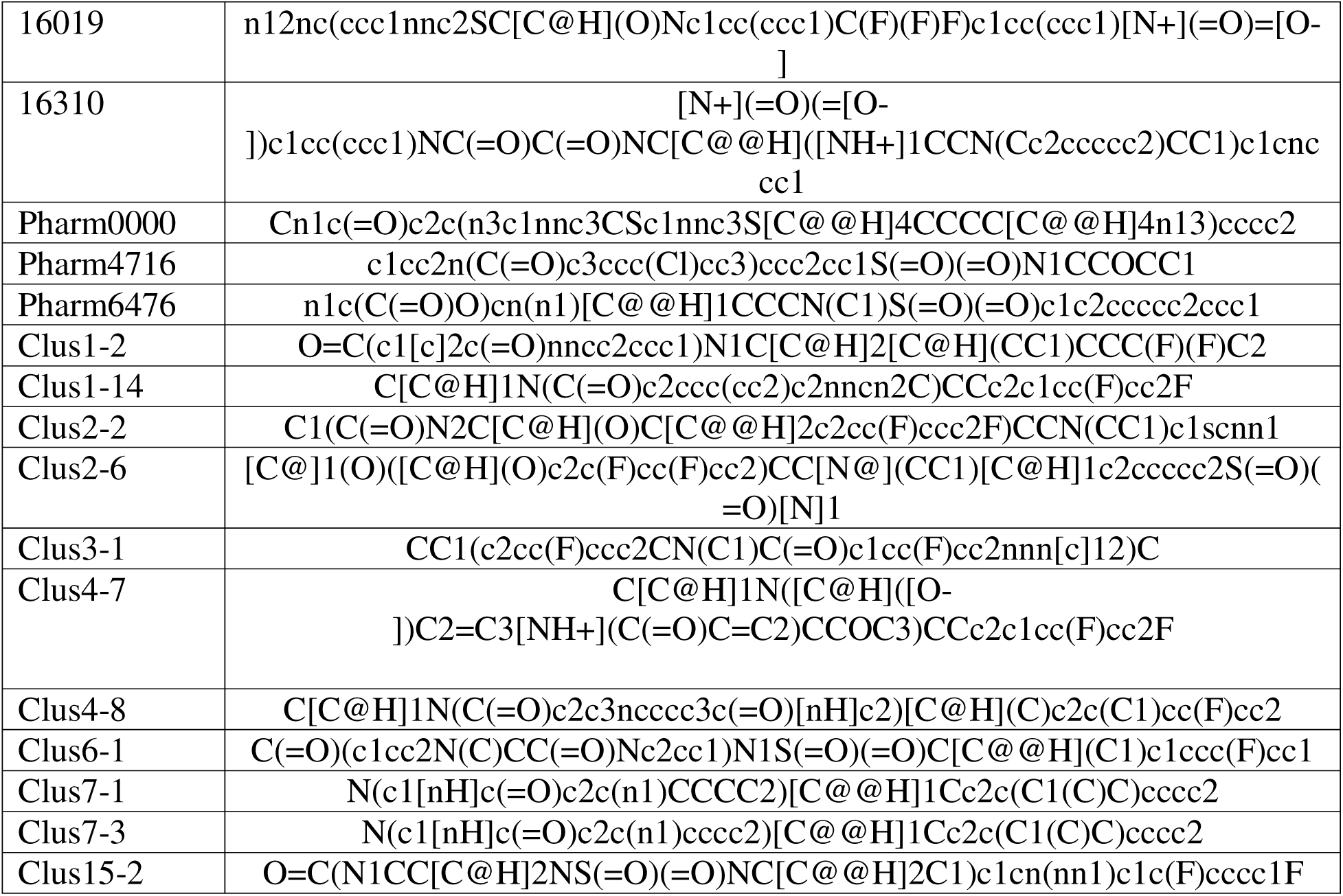
SMILES codes of studied compounds.

**Supplementary Table 3:**
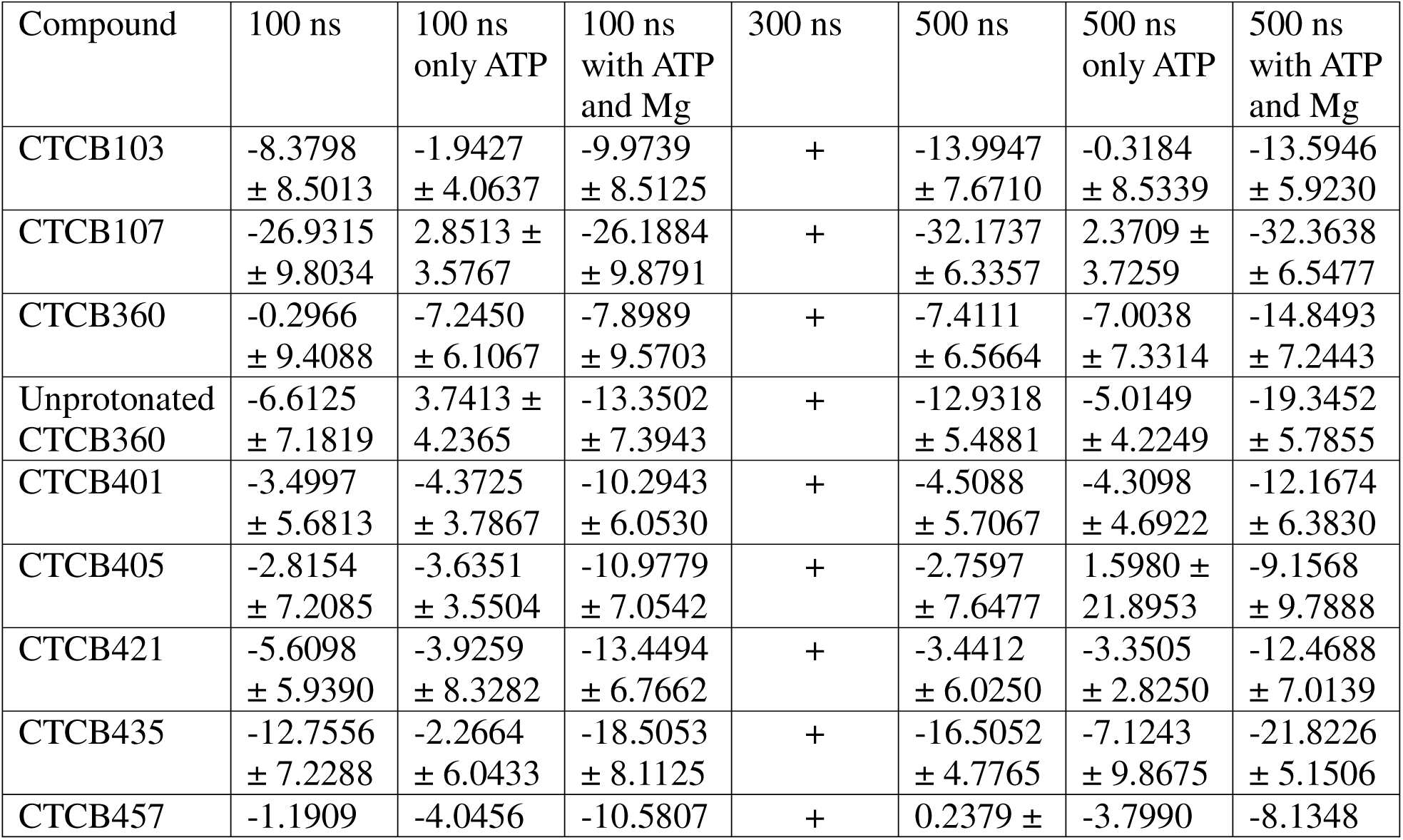
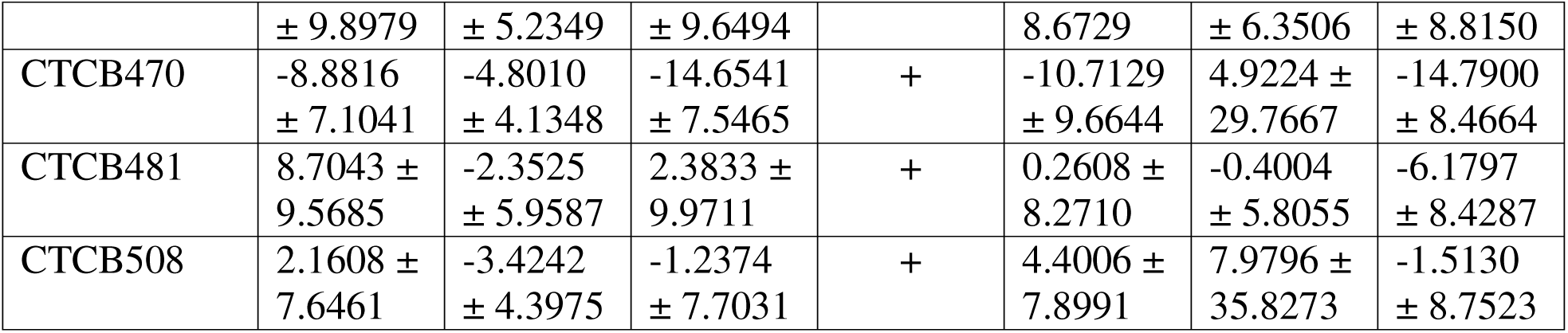
Results of MM-PBSA simulations for CTCB compounds.

**Supplementary Table 4:**
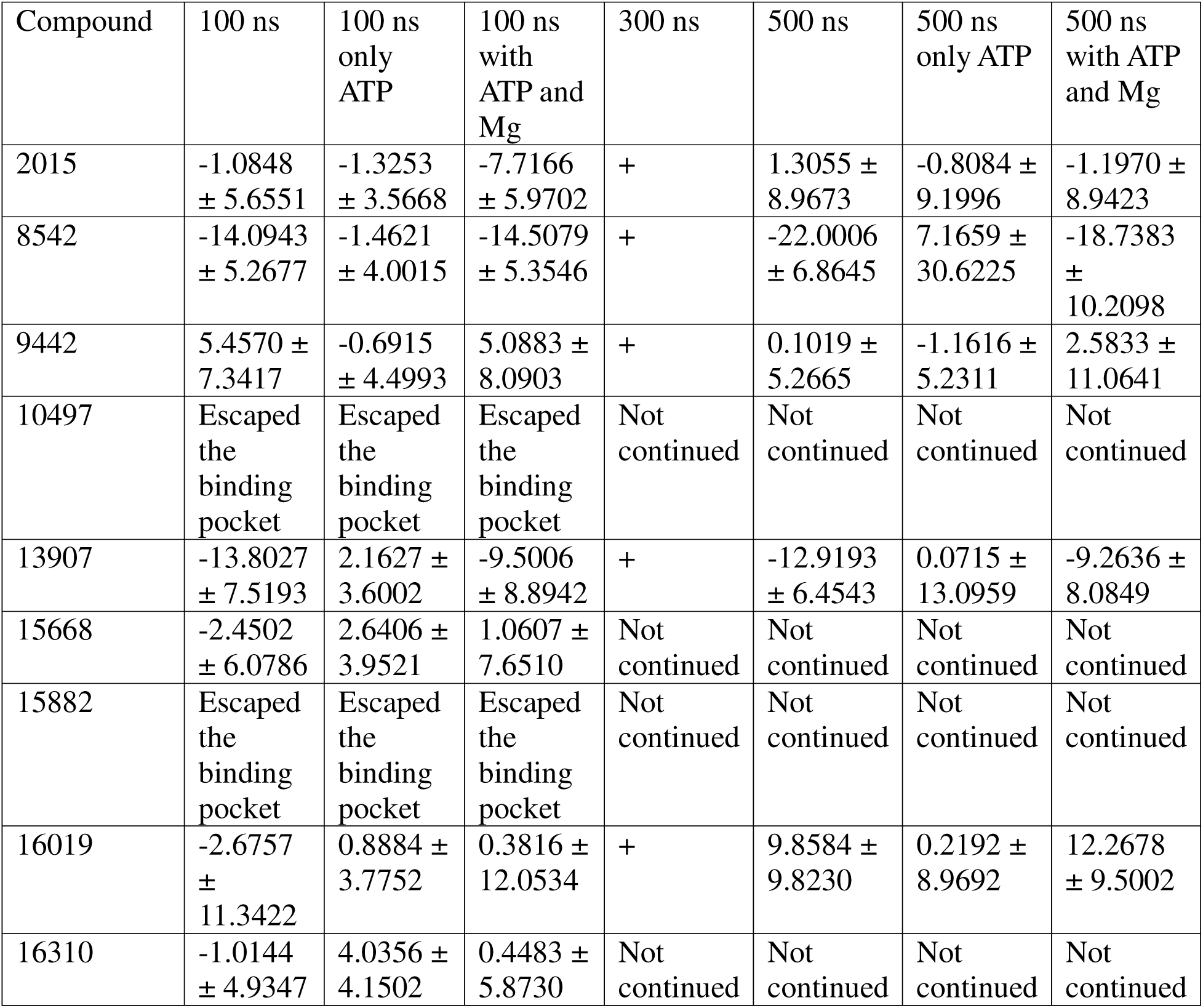
Results of MM-PBSA simulations for Life Chemicals kinase inhibitor library compounds.

**Supplementary Table 5:**
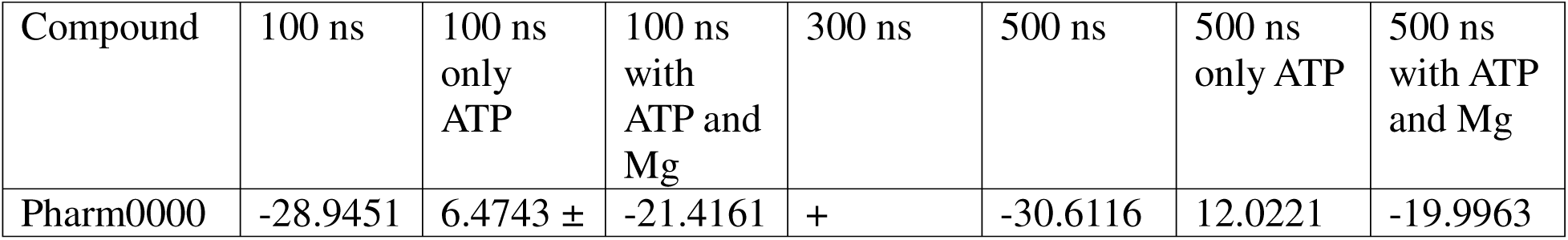
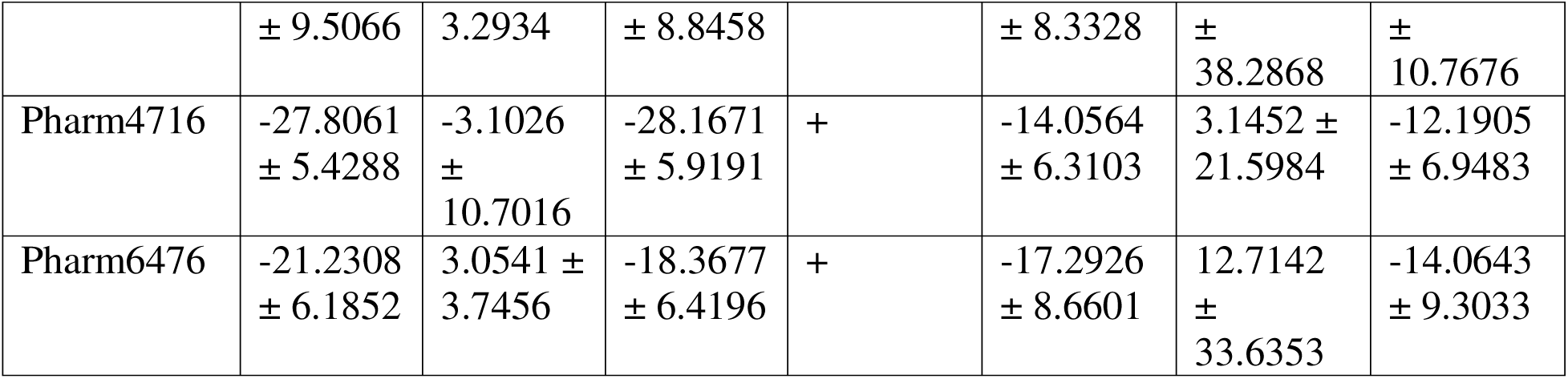
Results of MM-PBSA simulations for compounds derived from pharmacophore modelling.

**Supplementary Table 6:**
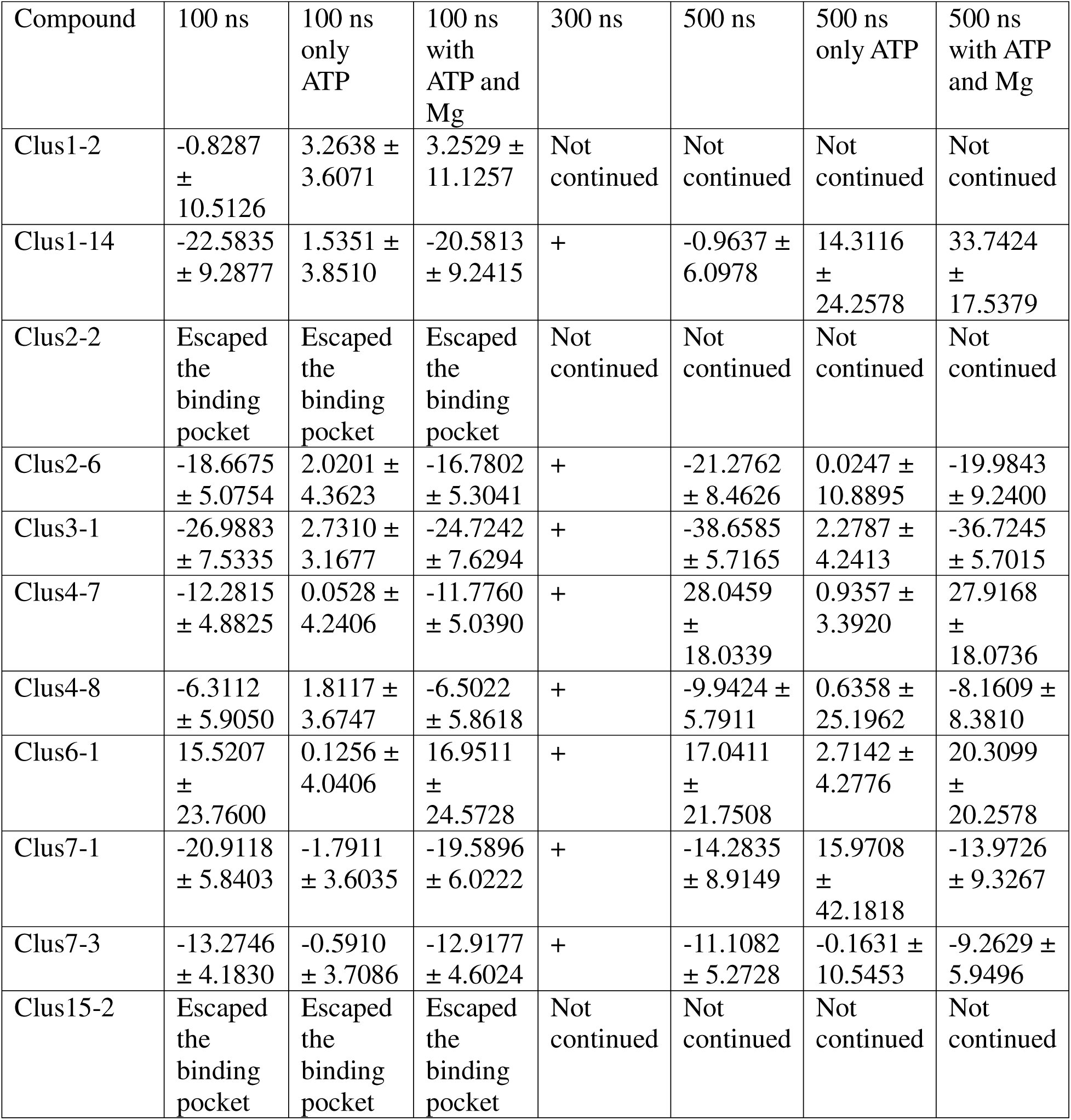
Results of MM-PBSA simulations for Enamine library virtual screen.

**Supplementary Table 7:**
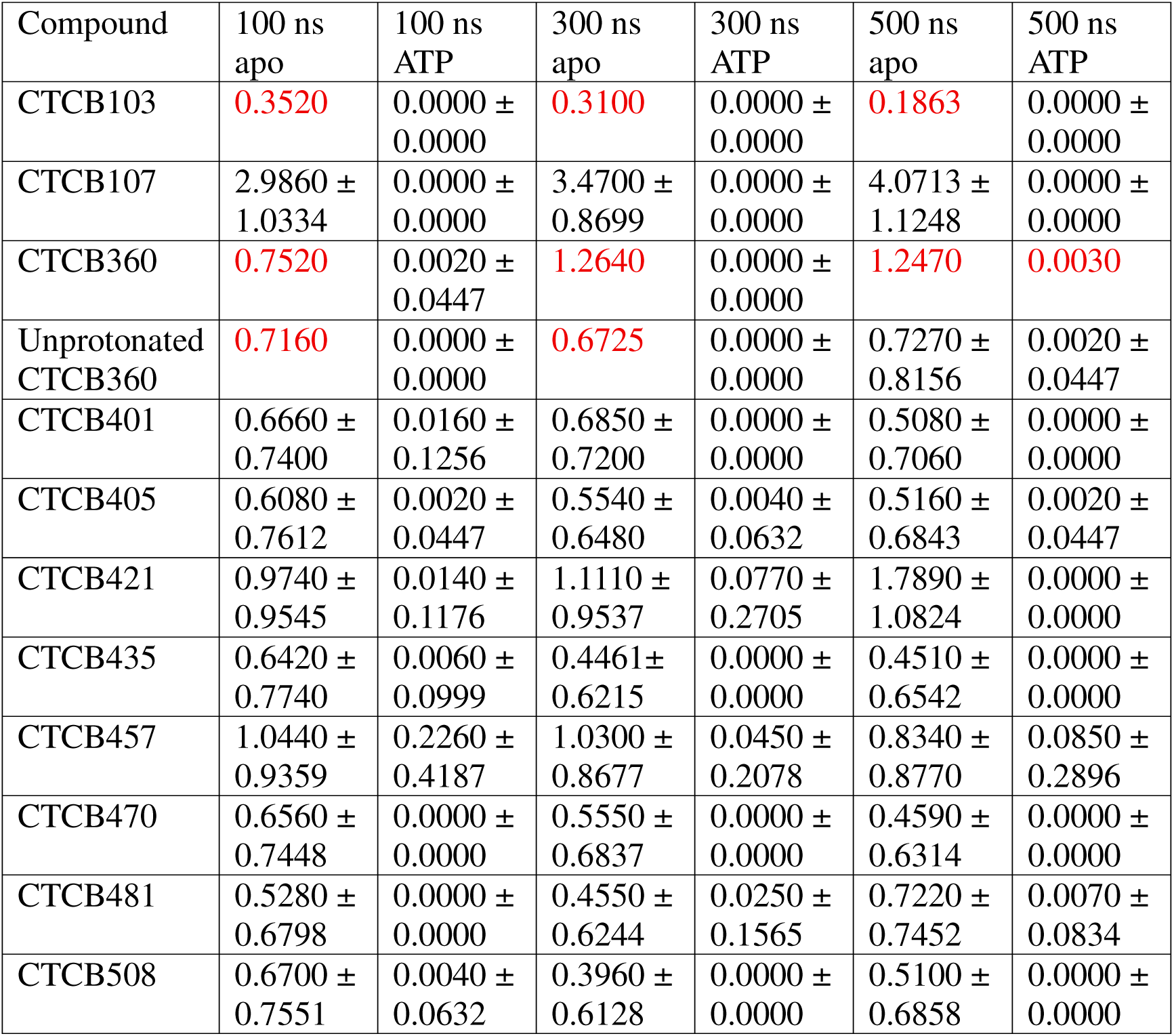
H-bond estimates for CTCB compounds.

**Supplementary Table 8:**
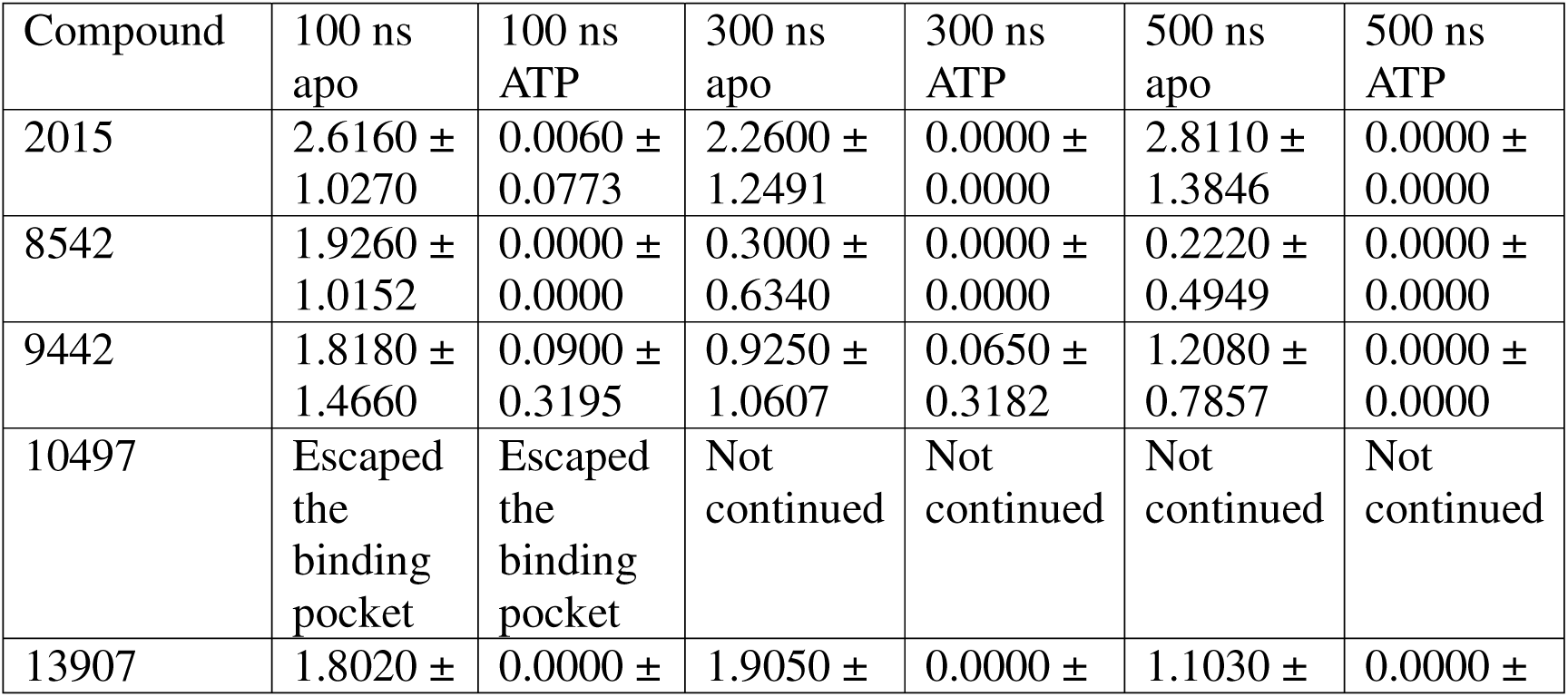
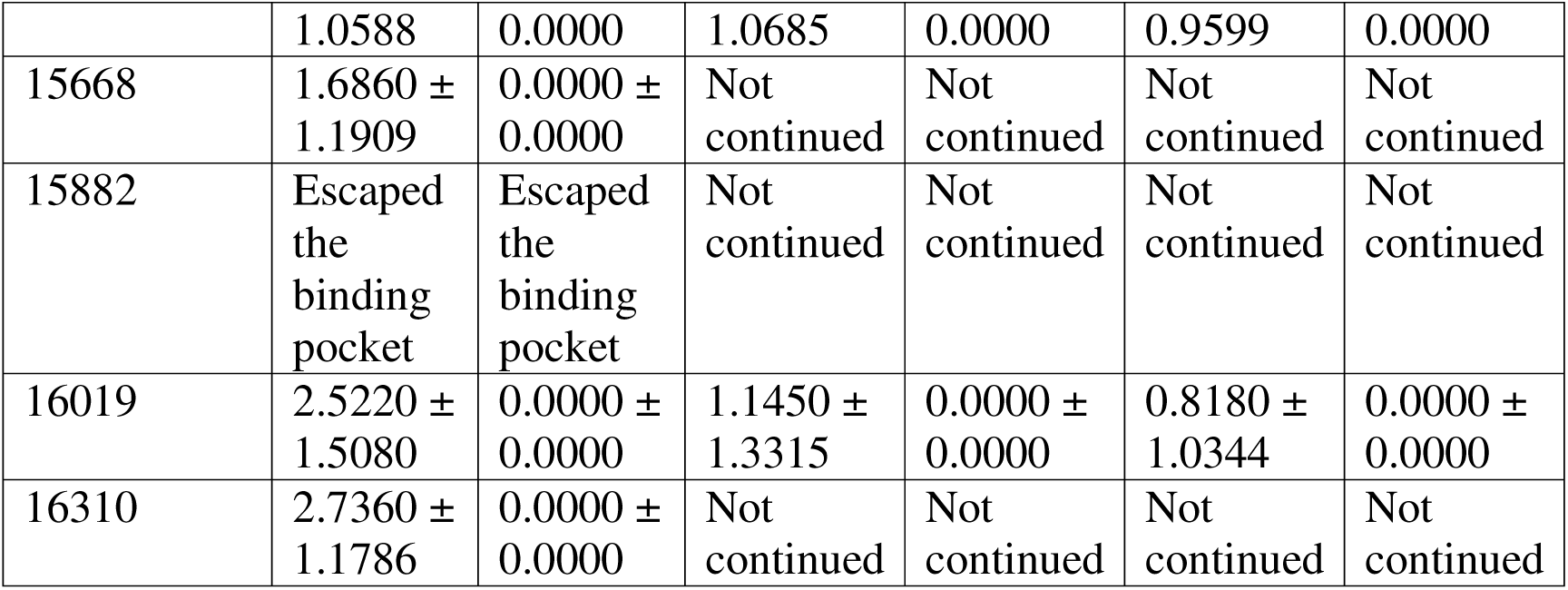
H-bond estimates for Life Chemicals kinase inhibitor library compounds.

**Supplementary Table 9:**
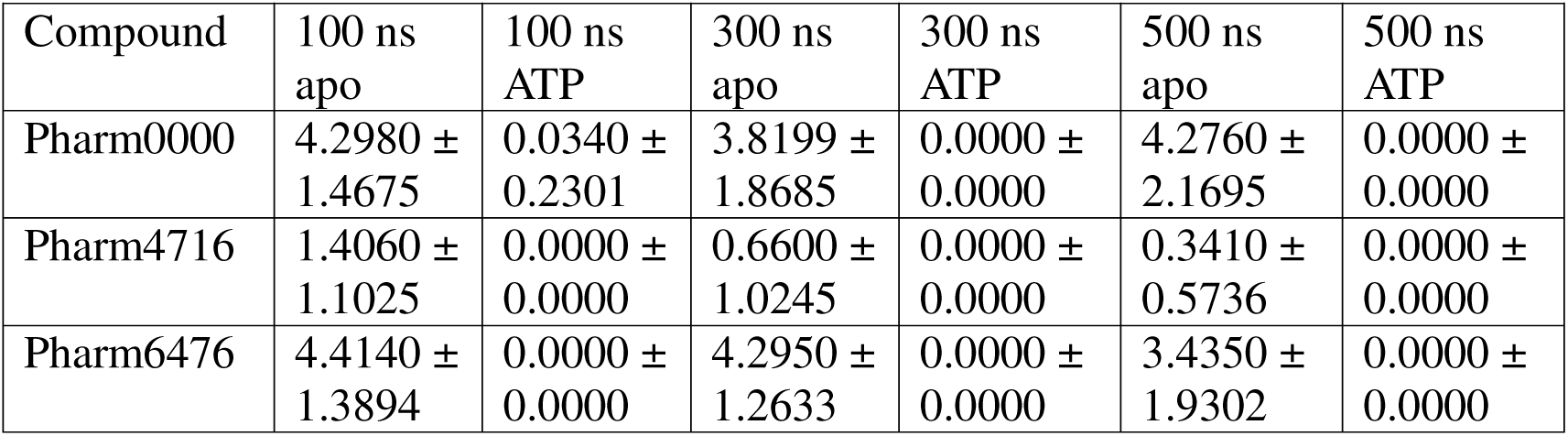
H-bond estimates for pharmacophore modelling-derived compounds.

**Supplementary Table 10:**
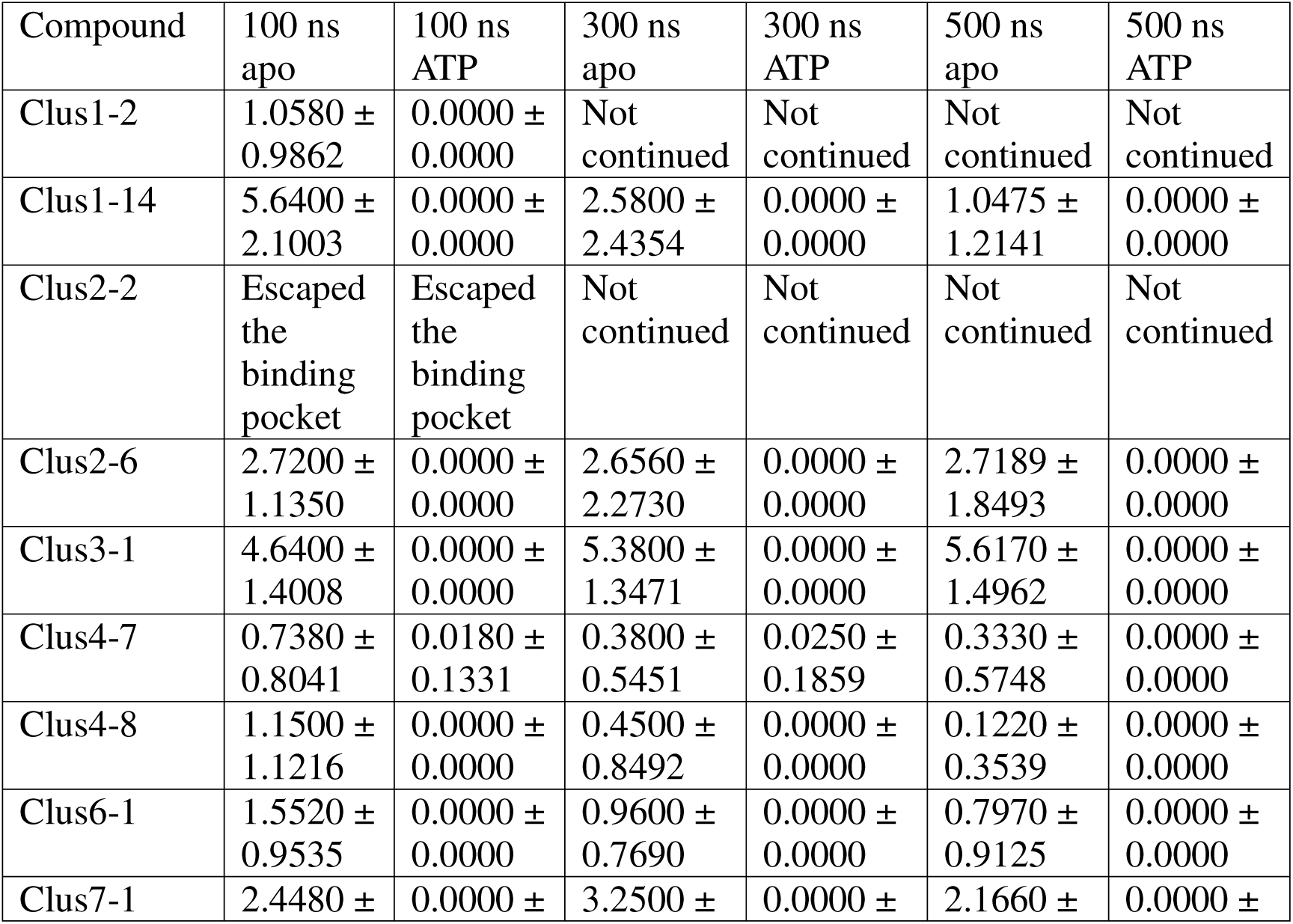
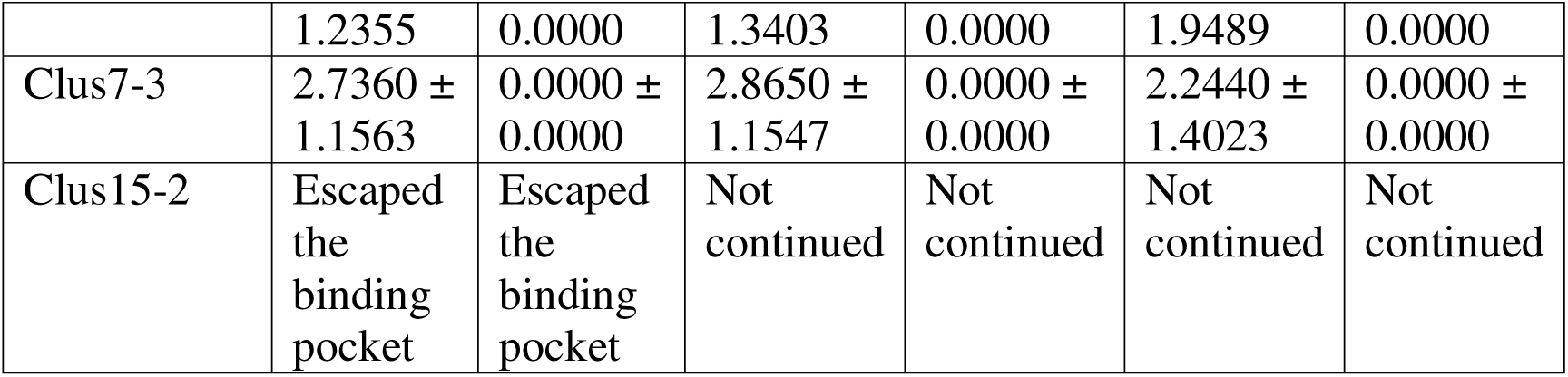
H-bond estimates for Enamine virtual screen.

**Supplementary Table 11:**
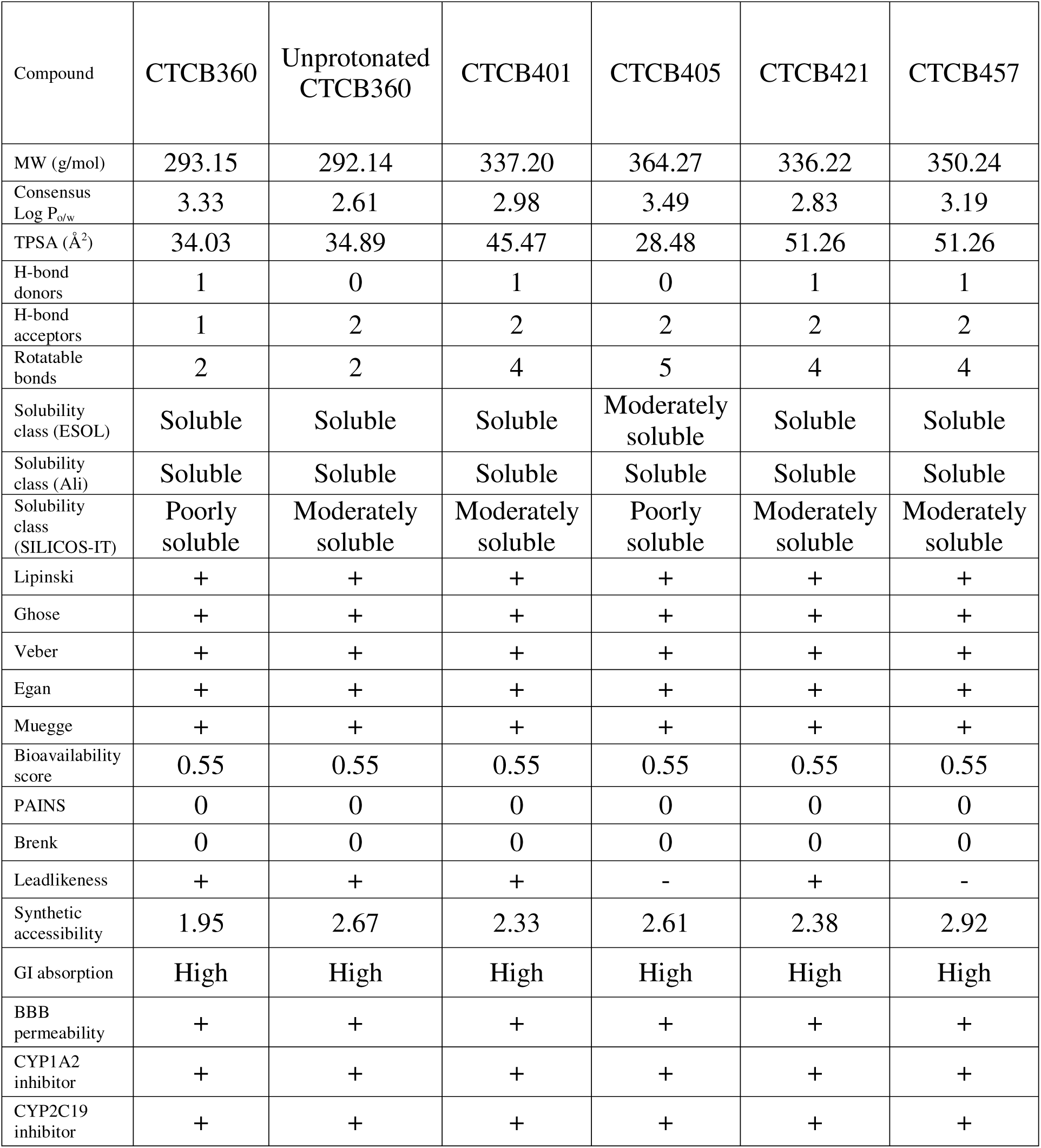
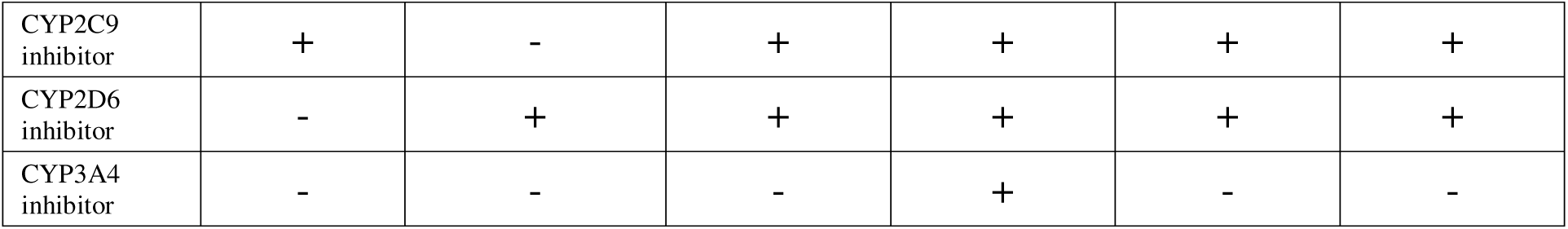
ADME prediction results for CTCB compounds.

**Supplementary Table 12:**
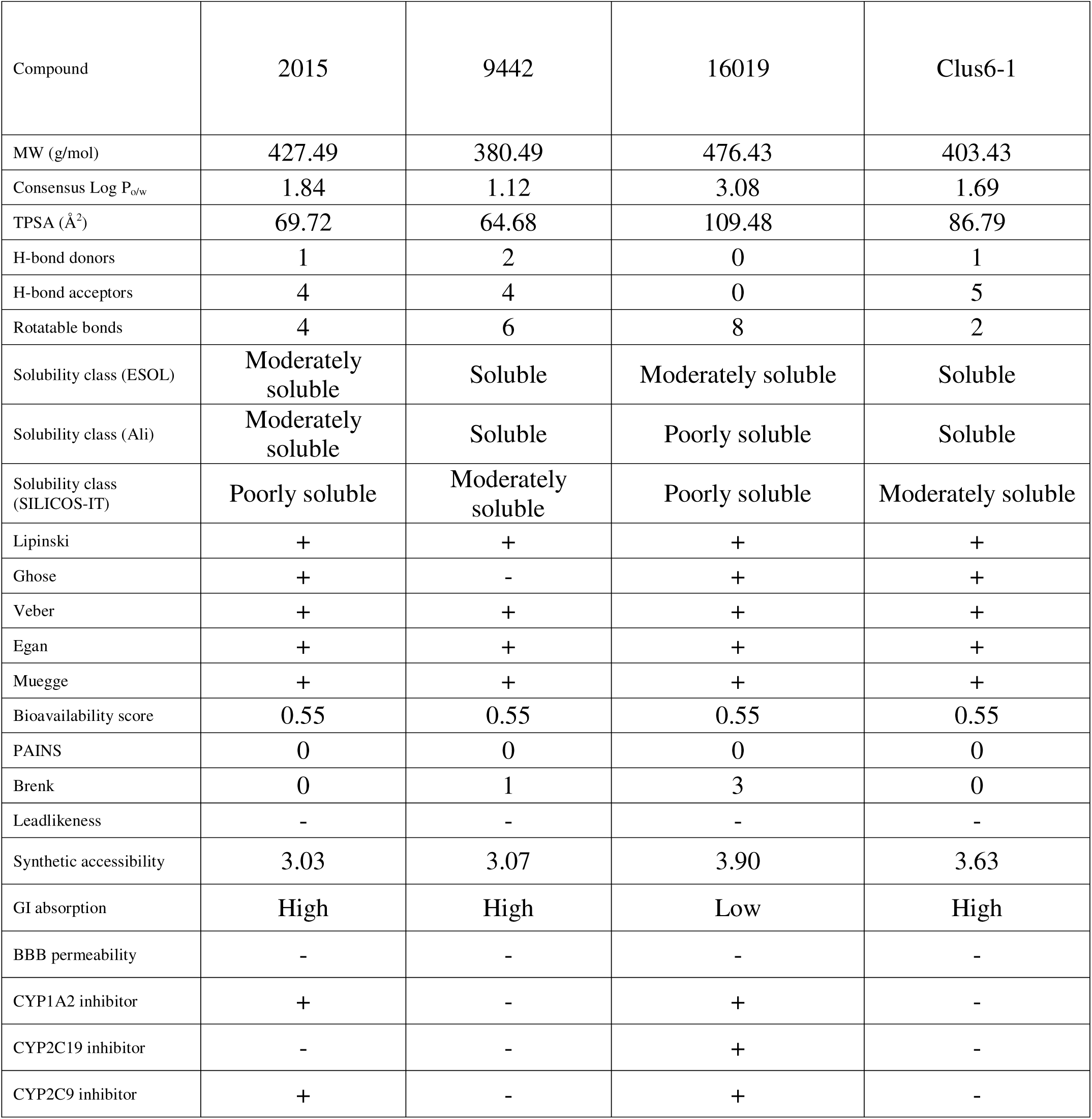
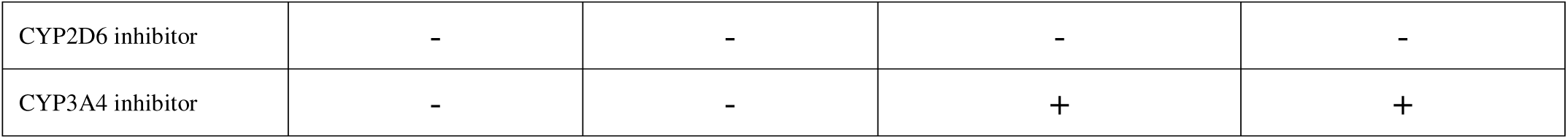
ADME prediction results for additional compounds.

**Supplementary Table 13:**
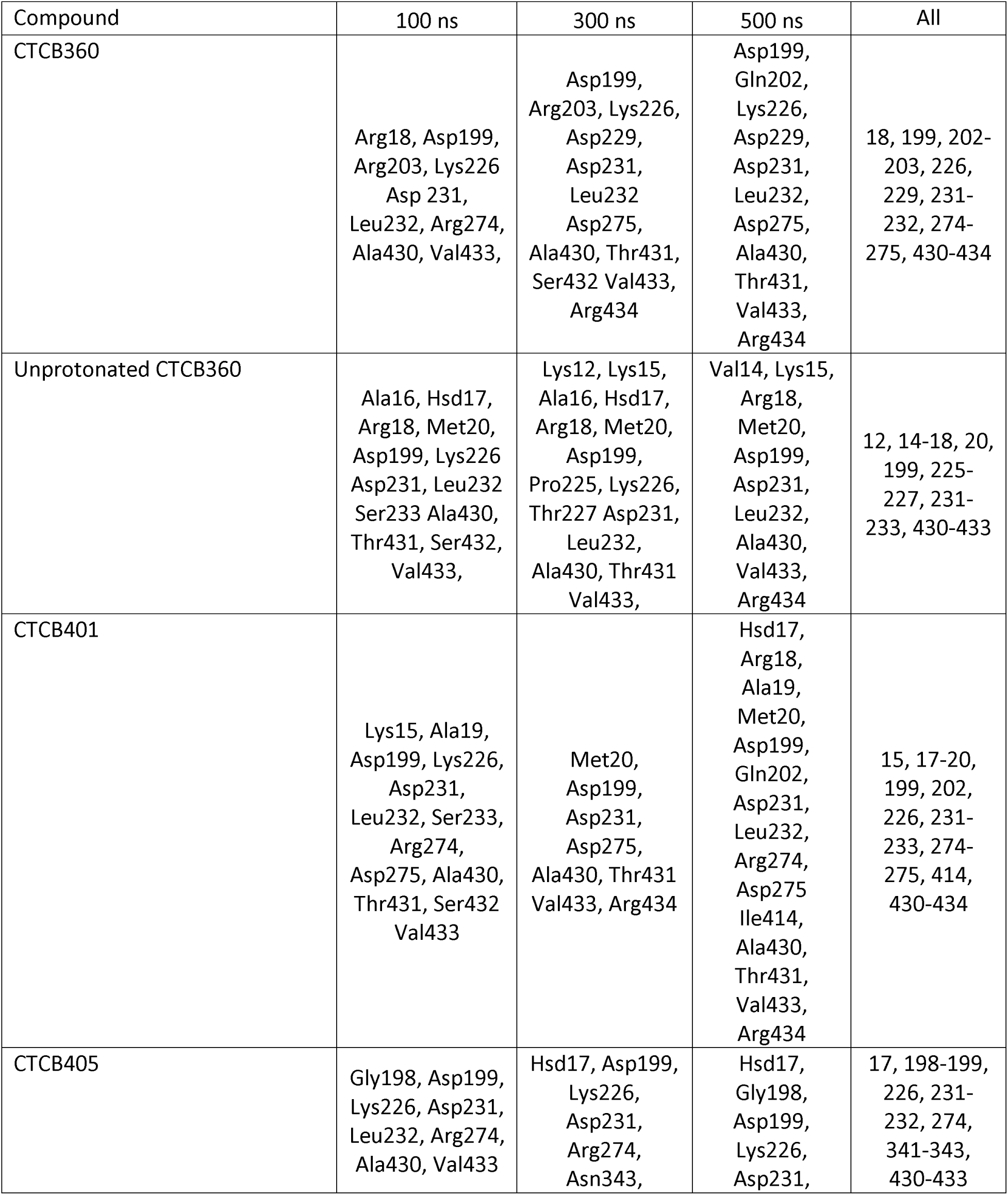
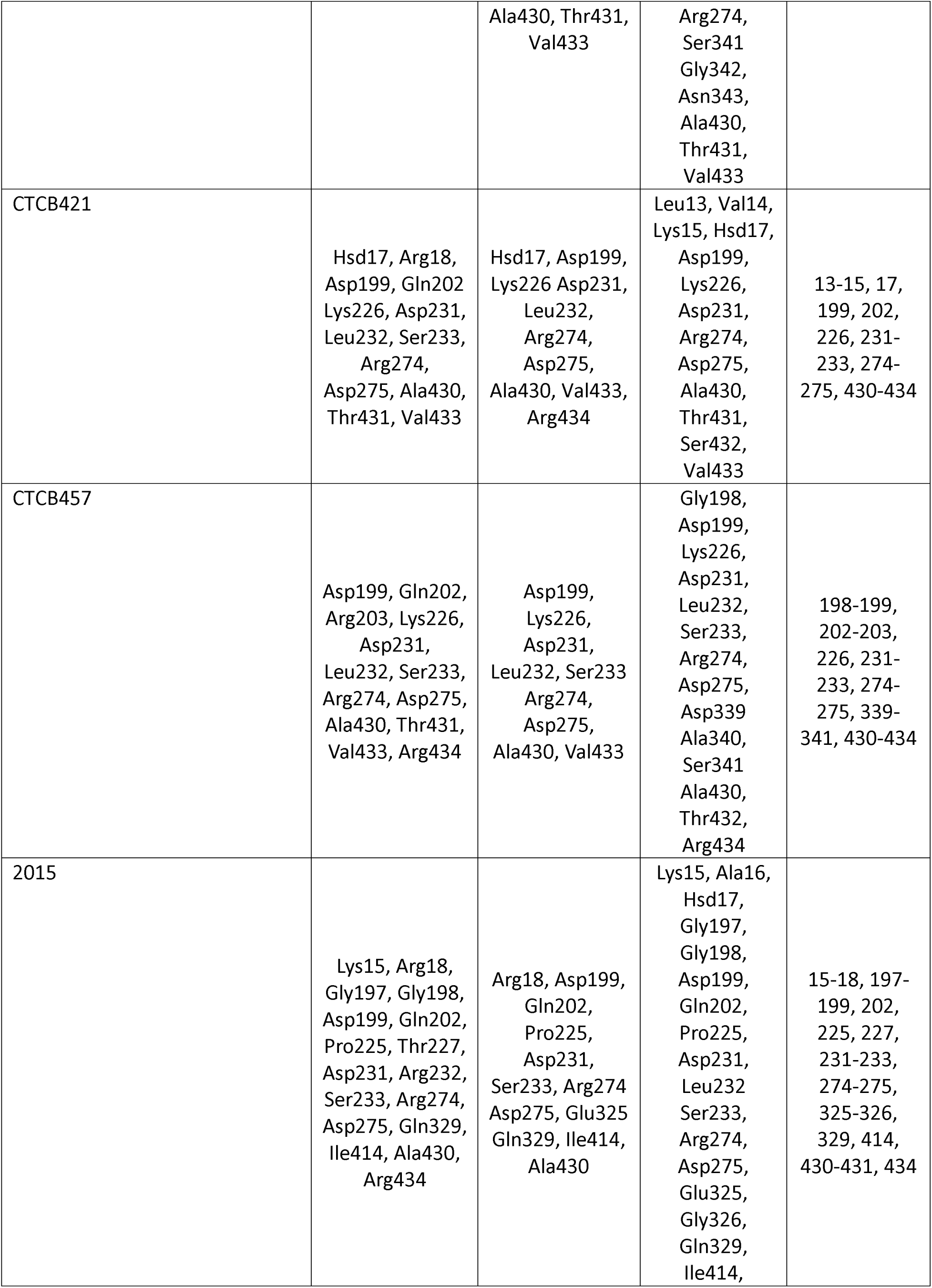
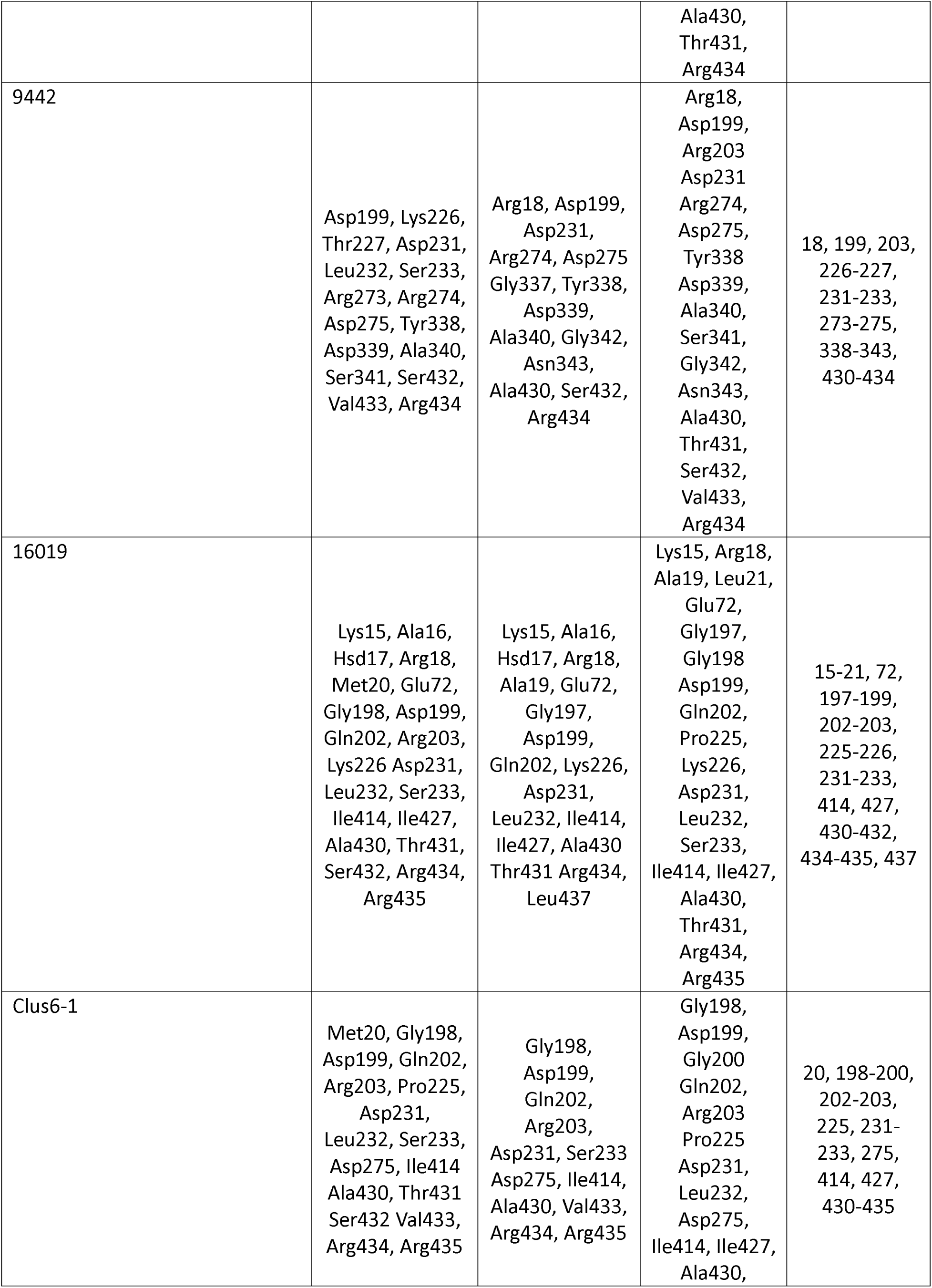
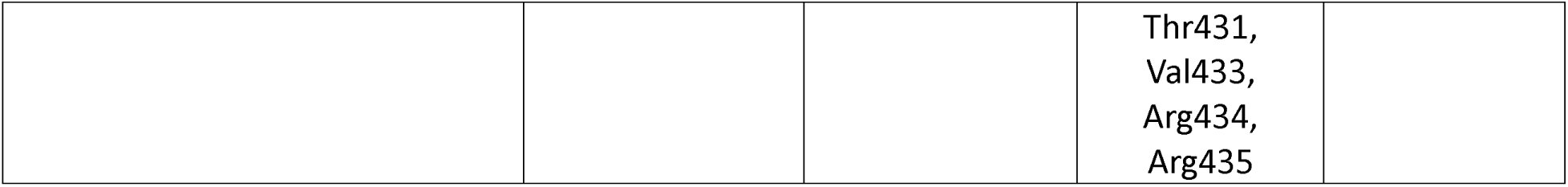
Hydrogen bond-forming residues for each of the selected compounds.

**Supplementary Table 14:**
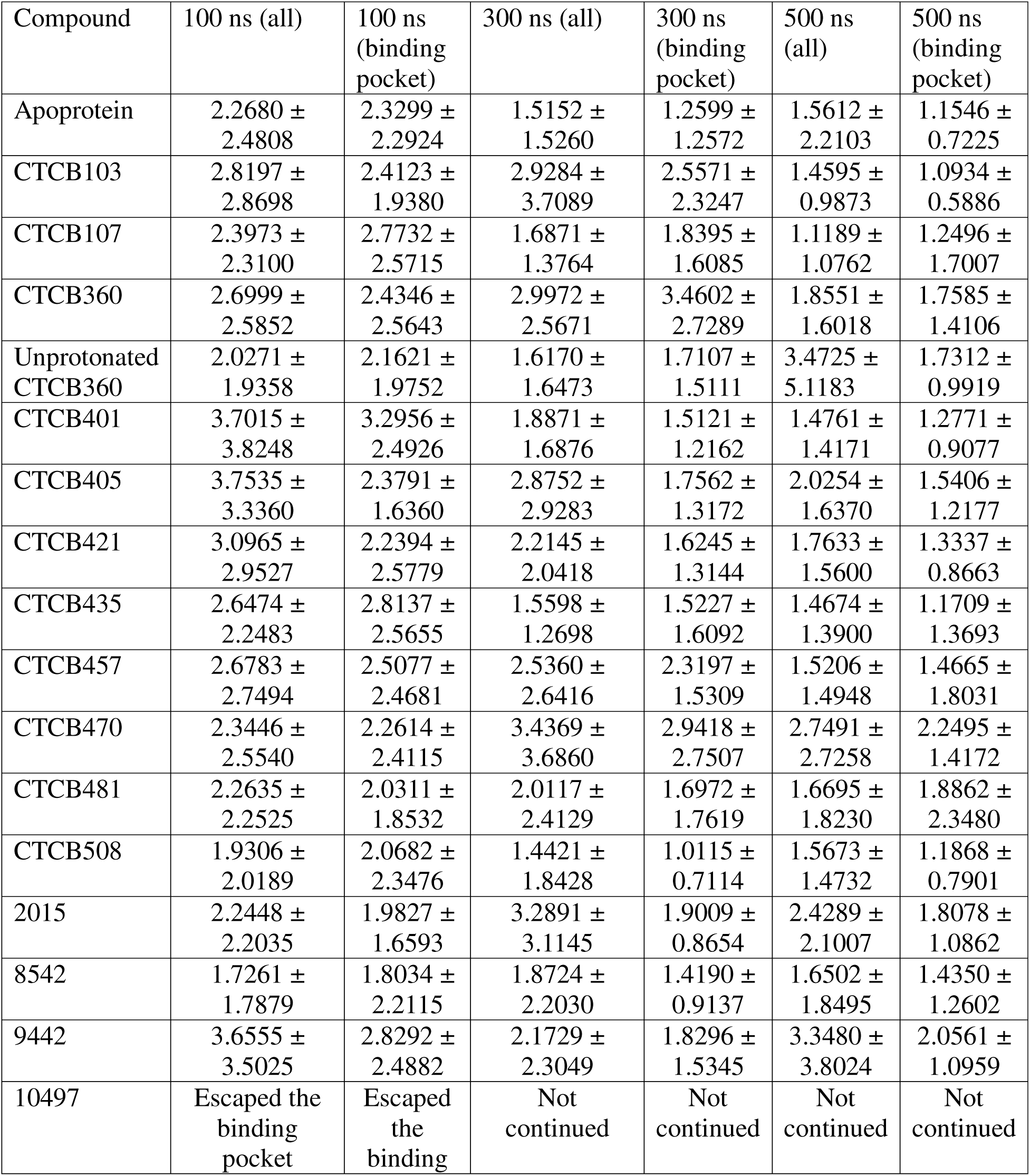
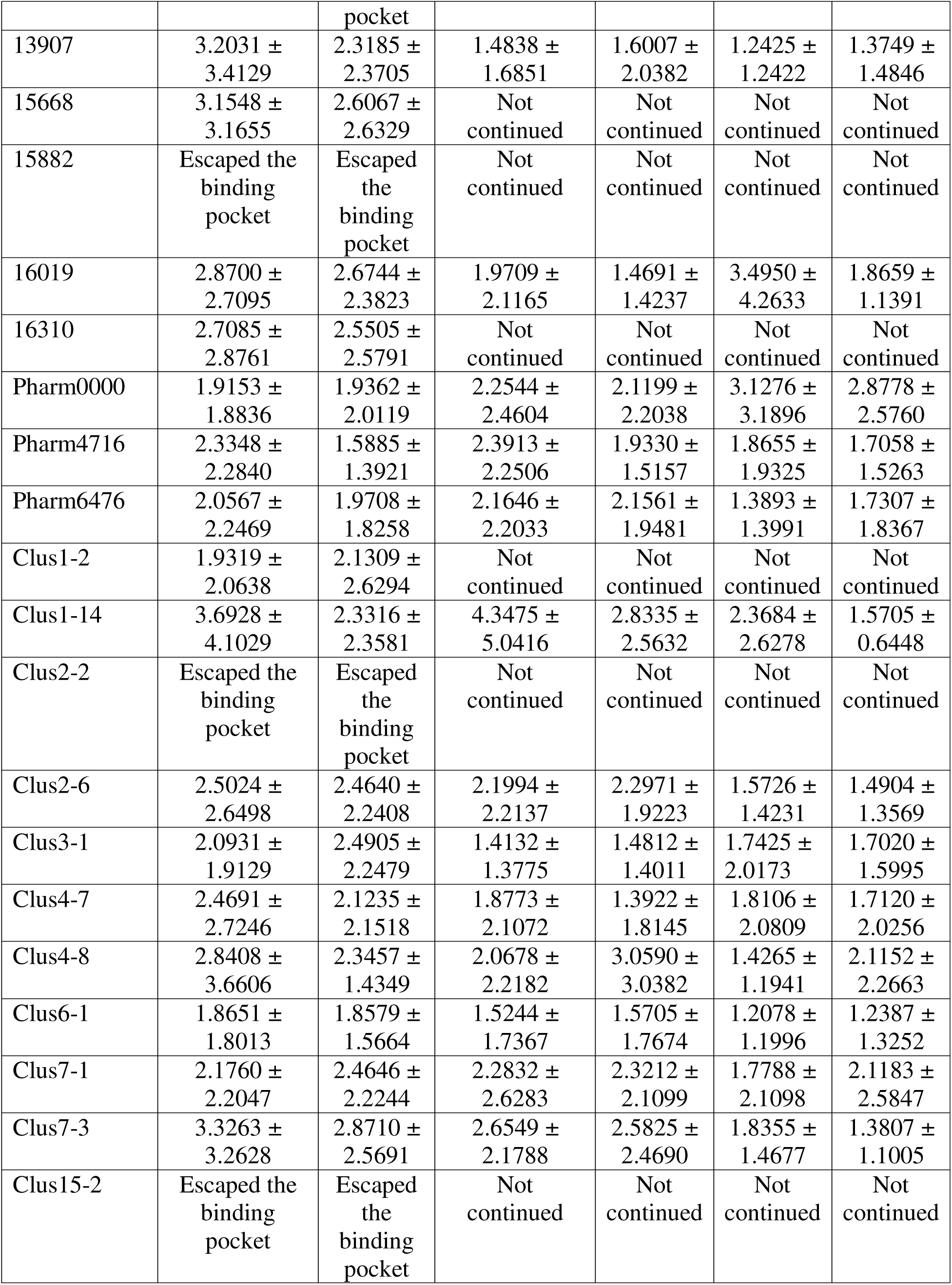
RMSF values for all residues and for the binding pocket.

## APPENDICES

### 1 Selection of Life Chemicals docking ligands

~~~
gnorm5pro <- read.csv("C:/Users/gedim/Desktop/gnorm5pro_full_scoretable.csv")
gnorm5pro_2 <- subset(gnorm5pro, gnorm5pro$Vina.SCORCH_score > 0.3)
gnorm5pro_3 <- subset(gnorm5pro_2, gnorm5pro_2$Vina.SCORCH_certainty > 0.5)
gnorm5pro_4 <- subset(gnorm5pro_3, gnorm5pro_3$AutoDock.SCORCH_score > 0.1)
for(x in 1:length(gnorm5pro_4$Ligand_ID))
gnorm5pro_4[[9]][[x]] <-
(gnorm5pro_4$Vina.Best.Score[[x]]+gnorm5pro_4$AutoDock.Best.Score[[x]])/2
gnorm5pro_5_o <- order(unlist(gnorm5pro_4$V9))
gnorm5pro_5 <- gnorm5pro_4[gnorm5pro_5_o,]
gnorm5pro_6 <- head(gnorm5pro_5, 3)
gnorm5 <- read.csv("C:/Users/gedim/Desktop/gnorm5_full_scoretable.csv")
gnorm5_2 <- subset(gnorm5, gnorm5$Vina.SCORCH_score > 0.3)
gnorm5_3 <- subset(gnorm5_2, gnorm5_2$Vina.SCORCH_certainty > 0.5)
gnorm5_4 <- subset(gnorm5_3, gnorm5_3$AutoDock.SCORCH_score > 0.1)
for(x in 1:length(gnorm5_4$Ligand_ID))
gnorm5_4[[9]][[x]] <-
(gnorm5_4$Vina.Best.Score[[x]]+gnorm5_4$AutoDock.Best.Score[[x]])/2
gnorm5_5_o <- order(unlist(gnorm5_4$V9))
gnorm5_5 <- gnorm5_4[gnorm5_5_o,]
gnorm5_6 <- head(gnorm5_5, 3)
f_6qu3 <- read.csv("C:/Users/gedim/Desktop/6qu3_full_scoretable.csv")
f_6qu3_2 <- subset(f_6qu3, f_6qu3$Vina.SCORCH_score > 0.3)
f_6qu3_3 <- subset(f_6qu3_2, f_6qu3_2$Vina.SCORCH_certainty > 0.5)
f_6qu3_4 <- subset(f_6qu3_3, f_6qu3_3$AutoDock.SCORCH_score > 0.1)
for(x in 1:length(f_6qu3_4$Ligand_ID))
f_6qu3_4[[9]][[x]] <- (f_6qu3_4$Vina.Best.Score[[x]]+f_6qu3_4$AutoDock.Best.Score[[x]])/2
f_6qu3_5_o <- order(unlist(f_6qu3_4$V9))
f_6qu3_5 <- f_6qu3_4[f_6qu3_5_o,]
f_6qu3_6 <- head(f_6qu3_5, 3)
~~~

